# BiomedCAT: Biomedical Context Analysis Tool for Supporting Mechanistic Hypothesis Generation via User-defined Contextual Subgraph Extraction

**DOI:** 10.64898/2026.11.11.766070

**Authors:** Sandrine Muller, Salah Eddine Chahma, Lucca Hoogenbosch, Laurent Torlay, Julie Dumonceaux, Christophe Hourdé

**Affiliations:** Univ. Grenoble Alpes, CNRS, Grenoble INP†, LIG, 38000 Grenoble, France; Grenoble INP† – Ensimag, MOSIG programme, Univ. Grenoble Alpes, Grenoble, France; NIHR Great Ormond Street Hospital Biomedical Research Centre, UCL Great Ormond Street Institute of Child Health, University College London, London, United Kingdom; Laboratoire Interuniversitaire de Biologie de la Motricité (LIBM, EA 7424), Université Savoie Mont Blanc, Chambéry, France

**Keywords:** Explainable AI, Biolink model, Knowledge graph, OCR, NLU, Contextual subgraph generation, Mechanistic hypotheses, FAIR

## Abstract

Researchers formulate mechanistic hypotheses in slides and figures long before they write them as text, yet knowledge-graph (KG) tools expect a query in the vocabulary of the graph. We present BiomedCAT, an open-source workflow that turns biomedical slides into a subgraph of the RTX-KG2c knowledge graph shaped by the user’s research profile, on a laptop and without sending the documents out. A small vision-language model describes each slide; a 4-billion-parameter agent types the terms into the classes of the Biolink model; an existence gate removes invented entities; identifiers are retrieved from Translator services or from a local table and selected, never generated, by a judge model; and a profile-weighted two-hop expansion builds the context graph. On a synthetic Biolink typing benchmark whose labels were audited by an expert, the local agent reaches 0.71 accuracy on 517 non-ambiguous mentions, 4 points below Claude Opus 5 (0.75), level with DeepSeek-V4-Flash (0.69) and above open-weight models of 24 and 31 billion parameters; the audit also showed that 55% of the generated labels were wrong or ambiguous. On expert-annotated slide decks, the reading stage recovers 81 to 94% of the explicit terms of the English deck, less on French decks, and linking assigns the expert identifier to 65% of the found terms online and 26% in the fully local mode. A one-slide run takes 33 to 42 minutes, with peaks of 9.8 GB of system RAM above idle and 5.3 GB of GPU memory. In a facioscapulohumeral muscular dystrophy (FSHD) case, three slides that each draw one pathway from DUX4 at a different biological scale give graphs that overlap by 7 to 15% of their nodes, each recovering its own pathway’s expected nodes, and expose sourced paths from DUX4, such as a DUX4–HMGB1 interaction from IntAct. Of 21 compounds tested in FSHD, seven, including every agent that targets DUX4, are absent from RTX-KG2c, which bounds what a graph-based approach can propose. BiomedCAT lets a researcher set a working hypothesis against curated knowledge, with every edge traceable to its source.

## 1 Introduction

The rapid expansion of biomedical literature has rendered manual curation unsustainable. PubMed alone receives more than 3,000 articles every day (Lee et al., 2019), outpacing the capacity of human annotation. Knowledge graphs (KGs) have emerged as the primary computational format for organizing biomedical knowledge, encoding concepts as entities with relationships as predicates to form a structured representation (Fang et al., 2023). However, entity representation and linking, and subgraph extraction are current limitations in making KGs accessible to non-experts. Resources such as PubMed Knowledge Graph 2.0, which covers over 36 million papers and over 482 million biomedical entity links, show the scale at which these links must stay unique and consistent across heterogeneous document sources. For downstream tasks, including drug discovery (Fang et al., 2023), biomedical data exploration and synthesis (Wang et al., 2026), and disease mechanism inference (Sudhahar et al., 2024), the reliability of entity extraction is the first requirement for the quality of link prediction and hypothesis inference (Xu et al., 2025).

Current approaches that open KGs to non-computational users mostly ground an LLM in the entire graph or in a large part of it (Edge et al., 2024). This introduces noise (Yu et al., 2025): the model may pull in biological facts that are connected in the graph but irrelevant to the question, while down-weighting the less established terms of the user’s query; the balance between the two is set by parameters that depend on the inference mechanism and are neither easy to customize nor to explain (Zhu et al., 2026).

Identifying the right biomedical entity is a prerequisite for link prediction and hypothesis generation (Zhang et al., 2025), and Named Entity Recognition (NER) on biomedical text adds a second layer of difficulty (Galea et al., 2018; Cariello et al., 2021; Kühnel and Fluck, 2022). Direct usage of general-purpose Large Language Models (LLMs) on biomedical text yields unsatisfactory results due to the shift between general and biomedical corpora, and general domain pre-training knowledge hinders models’ performance in the biomedical field (Gu et al., 2021). Fine-tuned domain-specific models outperform instruction-tuned LLMs on biomedical Natural Language Processing (NLP) (Fu et al., 2025), but the annotated corpora they are trained on follow the frequency of terms in the literature, which biases them towards well-studied areas. For instance, the widely used scispaCy models (Neumann et al., 2019) cover only a small part of the classes of the Biolink model and miss many finer-grained biomedical areas. These constraints matter most for hypothesis generation, which targets areas where knowledge is scarce, such as novel or rare diseases (Qin et al., 2025).

Moreover, the available pipelines take textual documents as input (Zhu et al., 2026) and are optimized for information retrieval rather than for fact-grounded hypothesis generation (Xu et al., 2025). Hypothesis inference is therefore grounded on either (1) a large corpus of published information containing mainly known and validated information; or (2) on a small set of human-curated queried named entities with user-independent rules (Sudhahar et al., 2024). However, in research settings, most of the information that could serve as a scientific context to create impact is highly dependent on tacit knowledge (Mitchell et al., 2022; Li et al., 2026). In User eXperience (UX) and Human–Computer Interaction (HCI) research, it is well established that the most valuable information for a user often lies in their immediate environment (tools, notes, visual aids) rather than in external databases, which led to contextual inquiry as a qualitative research method (Holtzblatt and Jones, 1993; Raven and Flanders, 1996; Ho et al., 2012) and to its integration with STEM technologies and AI (Hartmann et al., 2024; Lu et al., 2022; Cao et al., 2026). Likewise, what is needed to build novel and actionable hypotheses is often not in publications or patents, whose information value is protected, but in the tacit knowledge held in researchers’ presentations, a cognitive artefact of how they currently see a problem and judge how to solve it (Wedel, 2025; Miotto et al., 2018; Nedungadi et al., 2026). Reading images within KG retrieval pipelines, a modality that, to our knowledge, the KG retrieval literature has not addressed, could therefore bring these systems closer to the hypotheses researchers actually hold.

The third challenge is the normalization of entity identifiers, known as Biomedical Entity Linking (BioEL or BEL) (French and McInnes, 2023): the mapping of identified entities to their unique identifiers in a knowledge base (Kartchner et al., 2023), written in the Biolink model as compact URIs (CURIEs, such as MONDO:0001347) (Unni et al., 2022). Biomedical concepts carry large sets of names and aliases from different, non-unified knowledge bases. For instance, the same single gene can be referenced in multiple settings (Brown et al., 2015; Kinsella et al., 2011), and the correct identifier is context-dependent (e.g. taxon or sequence), which often requires specialized domain expertise for mapping accuracy (Kartchner et al., 2023). Directly generating identifiers with an LLM is unreliable, since models produce a plausible identifier rather than a verified one (Fu et al., 2025), a critical failure in a field where a well-formed but wrong identifier, such as that of a non-human orthologue, cannot be told apart from the right one without a lookup.

To our knowledge, no end-user system addresses these three challenges together. This paper introduces BiomedCAT, a transparent AI-assisted workflow that runs locally on an end-user computer and turns biomedical slides into Biolink-typed entities with their identifiers, and these into a user-defined context graph.

## 2 Materials and methods

BiomedCAT is an AI-assisted workflow for non-computational users that covers the stages required for an end user to get structured knowledge contextualized to their research hypothesis. It is organised in five stages (Figure 1):

- **Scientific description (reading)**: a vision-language model reads each slide image and writes a scientific description of it.
- **Biolink categorization (Named Entity Recognition, NER)**: the biomedical terms of the description are identified and typed into the classes of the Biolink model, a standard for biomedical knowledge representation, such as DUX4 as a gene and FSHD as a disease. Typing each term is what allows a precise linking to its known identifiers.
- **Verification**: an existence gate removes the typed terms that name no real entity; the user may then review the list.
- **Biomedical Entity Linking**: each entity is mapped to a standardized CURIE; for example, FSHD is resolved as MONDO:0001347. Candidate identifiers come from open Translator services (online mode) or from the local copy of the knowledge graph (fully local mode), and are canonicalised to the nodes of RTX-KG2c.
- **Context graph**: the linked identifiers seed a profile-weighted expansion of RTX-KG2c, a subgraph specific to the user’s reading of the field.

**Figure 1:**
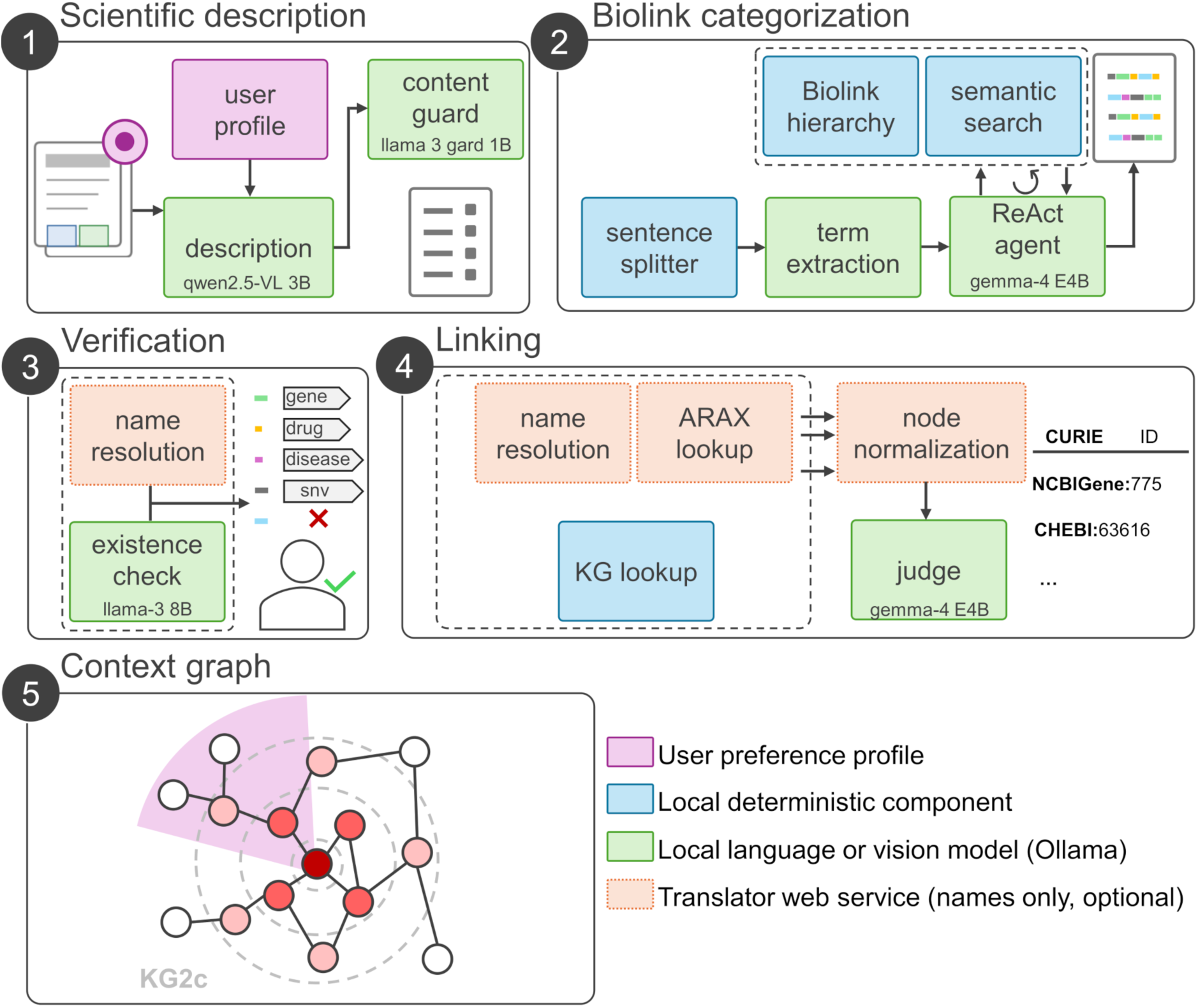
The BiomedCAT workflow. (1) A vision-language model describes each slide, guided by the focus area of the user profile and screened by a content guard. (2) Candidate terms are typed into Biolink classes by a ReAct agent querying three read-only tools over the Biolink model. (3) Typed terms are verified by a second model family and by the name resolver, with optional human review. (4) Candidate identifiers are retrieved, canonicalised to RTX-KG2c identifiers and selected by a judge model. (5) The identifiers seed a profile-weighted expansion of RTX-KG2c (rings: hops; shaded sector: profile scope). Green: local language or vision models; blue: local deterministic components; dashed orange: Translator web services, which receive entity names only and can be disabled; purple: user profile.

The application depends solely on small open-source models available in the Ollama library, chosen for their availability there and for their ability to run on a local machine with at most 16 GB of RAM and no GPU, the most common configuration among the end users we work with.

### 2.1 Base knowledge graph: representation of the scientific “world” knowledge

BiomedCAT uses the structured evidence of the RTX-KG2c knowledge graph (Wood et al., 2022) (KG; version kg2c-2.10.1-v1.0, aligned to Biolink Model 4.4.4) as its base knowledge of what is known in the literature. RTX-KG2c is a canonicalized integration of dozens of biomedical sources, in which equivalent identifiers are merged into one node. It holds 6,698,073 entities (nodes) and 26,948,303 relationships (edges); after the filtering described below, 3,097,290 nodes and 21,468,750 edges remain.

#### 2.1.1 KG filtering

We produced a filtered, compact copy that retains everything structurally usable by any down-stream analysis independently of a specific biomedical domain, allowing translational reasoning from the end user: they can ask domain-specific questions while keeping the information of connected domains. Five structural filters were applied, each targeting a source of noise:

- Non-Biolink model compliant nodes,
- Systematic curation of malformed edges,
- Equivalent mapping removal,
- Uninformative predicates, and
- Spurious associations through automated text-mining edges.

##### 2.1.1.1 Biolink Model alignment and malformed edges curation

Interoperability with the current Biolink Model was ensured by dropping predicates and node categories (i.e. entity types) that are not part of the current Biolink Model representation. A systematic repair of annotation artefacts was performed for the subset of predicates that carried a redundant doubled prefix (e.g. biolink:biolink_treats instead of biolink:treats); the 14,215 edges whose predicate is absent from the model were dropped.

##### 2.1.1.2 Equivalent mappings removal

Edges representing a near-equivalence between concepts (1,180,849 edges) were removed:

- biolink:same_as,
- biolink:close_match,
- biolink:exact_match,
- biolink:broad_match,
- biolink:narrow_match.

These links do not inform biomedical concepts interactions but rather near equivalent mapping between entities.

##### 2.1.1.3 Uninformative predicates and spurious associations

We excluded the predicates that are too generic to be informative (biolink:related_to, biolink:related_to_at_instance_level, biolink:related_to_at_concept_level; depth *d ≤* 2 in the Biolink predicate hierarchy), 2,617,669 edges.

##### 2.1.1.4 Noise from co-occurrence in literature and entity class mapping

We excluded the edges asserted only by automated text mining (occurs_together_in_literature_with, SemMedDB; 1,666,820 edges), so that the graph rests mainly on curated sources. UMLS nodes were excluded because our first typing tests revealed many generic nodes with the wrong class (e.g. modifier genes categorized as biolink:Gene), a source of spurious associations.

#### 2.1.2 Automated construction of scopes from end user profiles

BiomedCAT allows researchers to read the KG through different levels of granularity that we call profiles. Five profiles are shipped, each anchored on a few branches of the Biolink hierarchy: *biochemical actions* (molecular mechanisms: binding, interaction, regulation and pathway participation), *pharmacological intervention* (a chemical entity, its target, its mechanism of action and its clinical use), *clinical mechanisms* (disease, phenotype, predisposition, treatment and trial evidence), *epidemiological risk* (exposures, risk factors and disease occurrence in populations) and *genetic determinants* (the causal variant, its mode of inheritance and the gene or regulatory region it alters), together with a *uniform* baseline in which every branch and predicate has weight 1 (anchor classes and priorities: Supplementary Methods, *Profiles shipped with BiomedCAT*). The end user has the ability to build their own. A small set of human-defined anchor classes are bound to a scope. The scope is then expanded automatically through the Biolink model’s own class inheritance and grouping (i.e. mixin) relationships to produce different category scopes, each with its own vocabulary of relevant entity types and relationships.

Building on this structural representation, profiles were derived automatically from the scope within the Biolink class and predicate hierarchies. Researchers define their scope as a small number of high-level priorities over branches of the Biolink entity hierarchy: a branch is left out (*w* = 0), treated as secondary context (*w* = 0.5) or treated as primary (*w* = 1). From these few choices, the relevance of every applicable relationship and node category is derived automatically and reproducibly, and cached for reuse.

#### 2.1.3 Data store

##### 2.1.3.1 Biolink model information

For every Biolink class we built one text document holding its name, its definition and examples of usage, rewritten as natural-language sentences, which suit dense embeddings; the metadata (definition, examples, aliases, hierarchy) are also stored separately for exact and hierarchical search. Examples and aliases are truncated to five per class so that the context passed to the typing agent stays short.

##### 2.1.3.2 Exemplars addition

Class definitions are written in ontological language, whereas the text to be typed contains short surface forms (e.g. DM, type 2 diabetes mellitus). We therefore added to the index a set of exemplar mentions: 3,712 short expressions for 145 Biolink classes (25.6 per class on average, between 9 and 39), generated class by class with mistral-small in four surface forms: canonical names (1,174), multi-word compounds (1,023), abbreviations or symbols (774) and orthographic variants (741). Each exemplar is indexed as its own document carrying its class as metadata, and a query is matched against the closest exemplar; the resulting class ranking is fused with the definition-based retrieval by reciprocal rank fusion with the same weight as the dense leg. The exemplar strings also feed the exact lexical leg. Adding this leg improved the retriever hit@1 by 7 points (Supplementary Table S6, E1). A variant drawing the exemplars from the names and synonyms of RTX-KG2c nodes, stratified by class and weighted by node degree, was also built (Supplementary Methods) but did not improve retrieval (Supplementary Table S6, E2) and was not retained.

#### 2.1.4 Enrichment tables

Node descriptions, external identifiers, synonyms and edge citations were also extracted for display, together with the Gene Ontology (GO) annotations of the unfiltered graph, the universe of a hypergeometric GO enrichment.

### 2.2 Image processing through an OCR workflow

#### 2.2.1 Documents preprocessing

Documents are accepted as PDF files or as images (.png, .jpg, .jpeg), which avoids dependencies on proprietary software and preserves the spatial layout of scientific slides (tables, figures, text positioning) that is often lost when raw text layers are extracted; end users export their slides or schematics in either format, at 300 dpi for better recognition.

#### 2.2.2 Image description

Input slide decks are rasterized page by page and processed in small batches using qwen2.5vl:3b (Bai et al., 2025), a small open vision-language model designed for the understanding of complex document layouts with low memory use. In our tests it was the only model that interpreted the microscopy images, whereas the OCR-specialised GLM-OCR (Duan et al., 2026) transcribed text well but did not interpret figures (Section 2.5.1). The prompt asks the model for the detailed description that a translational biomedical scientist would give of the slide. The model is run at zero temperature so that the recall of relevant content is driven by prompt design rather than by sampling variability; in a factor test, sampling at temperature 1 added no content and corrupted entity names (Supplementary Methods, *Reading-stage factor test*; Supplementary Tables S4 and S5). Additional personalization is possible for specific entities on which the end user wants a particular focus (e.g. TP53). Pages are rasterized on demand and sent to the model four at a time, so that a long deck never holds all its images in memory.

#### 2.2.3 Input sanitization and personal information removal

Input text is cleared of common prompt-injection commands and screened by llama-guard3:1b for personal or offensive content before any other model reads it.

### 2.3 Biolink categorization from free text

The NER step determines the Biolink class of each term of the description. It addresses a practical difficulty: researchers rarely know how to express what they mean in a standardized, interoperable model. Typing is a retrieval-augmented classification performed by an agent over an index of the Biolink classes.

#### 2.3.1 Augmented processor

The Biolink model differs from traditional ontologies: rather than axiomatic descriptions of biomedical reality, it is engineered around the search and retrieval needs of subject-matter experts (SMEs), and its categorizations often differ from those of other ontologies, so the semantic proximity on which retrieval-augmented generation (RAG) relies does not guarantee proximity in the Biolink hierarchy. We therefore turned each class into a compact *learning object* (definition, ancestors, examples, aliases and exemplar mentions) suited to the tasks of the typing agent, in an index small enough for an end-user laptop. The agent, gemma4:e4b-it-qat with tool use and reasoning, explores the Biolink hierarchy and reasons, given the sentence, about the difference between the most probable classes for a term.

#### 2.3.2 Retrieval

The retrieval index is designed for low computational power: a light biomedical embedding model (bioclinical-modernbert-base-embeddings) encodes the class documents and the exemplar mentions into a local ChromaDB vector store, and a query is searched against both collections; the two rankings are fused by reciprocal rank fusion (Cormack et al., 2009) and the ten best classes are returned. Two further legs exist for exact surface forms, identifiers and compound terms: a sparse Okapi BM25 index (Robertson and Zaragoza, 2009) over spans tokenized with the en_core_sci_sm model, and an exact lexical match on class names, aliases, exemplars and KG2c synonyms. Both can be enabled by configuration; they are switched off in the published runs: the BM25 leg brought no gain, and the exact lexical leg, although it raised retriever hit@1 on the mentions of a real deck, left the accuracy of the agent unchanged (Supplementary Table S6, E11 and E12).

#### 2.3.3 Agentic NER

Many terms have a different meaning depending on their context; we call them *ambiguous* terms. For example, *human insulin resistance* can be a:

- biolink:PathologicalProcess: “a biologic function or a process having an abnormal or deleterious effect at the subcellular, cellular, multicellular, or organismal level” (Unni et al., 2022), as in *“Insulin resistance is a pathological process very frequently associated with NAFLD [. . .]”* (Memaj and Jornayvaz, 2022);
- biolink:PhenotypicFeature: “a combination of entity and quality that makes up a phenotyping statement; an observable characteristic of an individual resulting from the interaction of its genotype with its molecular and physical environment” (Unni et al., 2022), as when insulin resistance is reported as a trait of a patient.

For these reasons, typing a term from its context allows scientists to correctly identify their vocabulary of interest thereafter through linking, and not associate a broad concept to an intended specific meaning.

The NER process is executed through a Reason-Act-Observe (ReAct) agentic framework with the quantized gemma4:e4b-it-qat model, in three stages.

##### 2.3.3.1 Stage 1: Term Extraction (High Recall)

The first module is a high-recall extractor: the LLM is prompted to list, as a JSON array, every biomedical term of the sentence given as context, without filtering. Multi-word candidates are kept intact when they form a single semantic unit to preserve their meaning (e.g. *CACNA1C transcript* is different from *CACNA1C protein* where *CACNA1C* refers to the gene concept and its products are either *transcript* or *protein*). Purely numeric or non-alphabetic tokens are discarded.

##### 2.3.3.2 Stage 2: Entity Classification (Precision) through agentic reasoning

Each extracted term is then passed to a second module, which assigns it the single most relevant Biolink class with the help of the retrieval index. The agent follows a ReAct strategy: at each reasoning iteration, it may either call one of three read-only tools (semantic search over Biolink class hierarchy, exact term lookup, and Biolink class neighborhood search) or commit to final verdict. Semantic search queries the retrieval index described above, whose class documents include the ancestor chain. This lets the agent ground its decision in the Biolink class closest in meaning to the term rather than in its surface form; it has to reason on the adequacy between the meaning of the term and the class definition, and otherwise fall back on the neighbouring classes and iterate.

##### 2.3.3.3 Stage 3: Validation and sensitive data gating

Typing models invent entities from spelling: a plausible-looking symbol receives a class whether or not it exists. An existence gate is therefore applied once per document, after the typing model has been released from memory, to the list of typed terms. Two independent checks are run on the bare term, without its class or its sentence. The first asks the Translator Name Resolver from Babel (Morris et al., 2026) (production instance, v1.7.0, Babel release 2026jul22) whether the string names anything at all, in any Biolink category; a network failure keeps the term, so that the gate never rejects on connectivity. The second presents the terms, in batches of forty, to a model of another family than the typing model (llama3:8b) prompted as a strict biomedical fact checker that lists the strings that do not name a real biomedical entity. In the published configuration (union) a term is dropped when either check fails; the modes nameres, llm and off are available for ablation. In the fully local mode no term may leave the machine, so the resolver check is skipped and only the second-family model remains. Terms that later receive no identifier at the linking stage are kept with an explicit NIL: the gate decides existence, not linkability. Mentions are then deduplicated case-sensitively, so that biologically distinct entities (Co for cobalt, CO for carbon monoxide) remain distinct while repeated extractions of the same term collapse. Personal data is handled upstream and downstream of the gate: the typing agent answers NONE for identifiable people, and terms typed into person-denoting classes (case, individual organism, cohort, study population, agent) or matching identifier patterns (dates, e-mail addresses, phone and record numbers) are never sent to an external service.

#### 2.3.4 Entity linking

Typed terms are linked to an identifier in two selectable modes. In the online mode a candidate pool is assembled for each term from three lookups that return names only: the Translator Name Resolver without type constraint, the ARAX entity endpoint (Glen et al., 2023), and one further Name Resolver query per Biolink class predicted for the term, constrained to that class (biolink_type), so that the class decided at stage 2 orients the candidates without excluding those of the unconstrained pass. Every candidate identifier is mapped to its canonical node in the local RTX-KG2c equivalence table; identifiers absent from that table are first sent to the Translator Node Normalizer (conflation off) and its preferred clique identifier is looked up in turn, so that an identifier returned under another prefix still reaches the graph node when one exists. The pooled candidates, each shown with its label, its Biolink category and whether it is present in KG2c, are presented to the typing model acting as a judge (*LLM-as-a-judge*), which selects one candidate or abstains; the selected identifier and its KG2c canonical node are both recorded. In the fully local mode no term leaves the machine: candidates come from an exact match of the term against the names and synonyms of the local KG2c node table, restricted to the predicted class. The versions of the remote services at the time of the published runs are recorded with the runs (Name Resolver v1.7.0 on Babel 2026jul22 and Biolink model 4.4.3; Node Normalizer 2.3.18; ARAX API v1.4), and the Biolink model used by the pipeline is pinned to release 4.4.4.

#### 2.3.5 Empirical selection of the typing configuration through iterative evaluation

The configuration of the typing step was not fixed a priori but selected through twenty-one single-change experiments (Supplementary Table S6), each stated as an observation, a hypothesis and an acceptance rule defined before the run. Retriever changes were scored by hit@1 on the 1,075 public synthetic terms, and agent changes by accuracy on a fixed 300-term reference subset. A change was adopted only if it improved the score by more than 5 points, a margin larger than both the sampling noise of the subset (about 2.8 points) and the run-to-run variation observed on the full benchmark (about 1 point). Because the subset was scored against the original LLM-generated labels, which the expert audit later showed to be partly wrong or ambiguous (Section 2.5.2.3), only differences between configurations are interpreted.

### 2.4 Context graph construction

The last stage turns the linked identifiers of a document into a subgraph of the filtered RTX-KG2c that is specific to the user’s reading of the field. It is a profile-weighted best-path expansion executed as SQL over the KG2c Parquet tables (DuckDB, biomedcat/stages/context_graph.py); no model is involved, so the same seeds and the same profile give the same node scores. Two builds can differ only in which nodes of equal score fill the last places of a saturated category, an order that the database engine leaves free: two builds of the FSHD graph differ by 86 of their 5,878 kept nodes, all tied at the cap.

#### 2.4.1 Seeds

Every typed mention that received a canonical RTX-KG2c identifier at the linking stage becomes a seed; the mentions and slides it comes from are kept as provenance, so that a node of the final graph can be traced back to the slide that introduced it. Seeds whose identifier is absent from the filtered graph are reported and skipped.

#### 2.4.2 Edge weighting

Each edge of the filtered graph receives a weight from the active profile: the product of its predicate weight, its primary knowledge-source weight and its knowledge-level weight, as derived from the user’s branch priorities (Section 2.1.2). Edges whose predicate has weight zero in the profile, or whose weight falls below a minimum, are not traversed; edges asserted as predictions can be excluded. The profile also defines a class scope: an edge whose endpoint belongs to a Biolink category outside the scope is dropped, unless that endpoint is a seed, so that out-of-scope seeds remain reachable. Representation edges, which join a gene to its protein and to its drug-target identity, carry predicate weight one whatever the profile, because crossing between the identifiers of the same biological object is not a mechanistic step. Traversal is directed by default: an edge is followed from subject to object at its full weight and from object to subject at a fraction of it (0.5 in the published profiles), except for symmetric predicates such as *interacts with*, which are never penalised; profiles may switch the traversal to undirected.

#### 2.4.3 Expansion

Starting from the seeds at hop 0, the expansion proceeds hop by hop (two hops in the published runs). At each hop the frontier is the set of nodes reached at the previous hop; the weight of every path through a frontier node is divided by log_2_(2 + deg(*u*)), where deg(*u*) is the undirected degree of the node in the filtered graph, so that a path through a generic hub (a disease like cancer, a housekeeping protein) contributes little; seeds themselves are not penalised, and nodes above a hub cap of 500 edges are reached but never expanded. The frontier is joined with the weighted edge table and, for every pair (seed, node), only the best path weight is kept, the product of the edge weights along the path divided by the degree penalties of its intermediate nodes; a node already reached from a seed at an earlier hop keeps that hop. After each hop, a bounded closure (two steps) extends the newly reached nodes along representation edges without consuming a hop, so that reaching a gene also reaches its protein at the same distance.

#### 2.4.4 Scoring and pruning

Every reached non-seed node *v* is scored as the sum over seeds of its best path weight, divided by log_2_(2 + deg(*v*))*^α^* with *α* = 0.5, a milder penalty on the target node’s own degree; the number of seeds that reach the node (its coverage) and its hop are recorded with it. Two prunings follow. A retention floor removes, within each hop separately, the nodes below the median score of that hop, because the score scales differently at each hop and a single threshold would act on one hop only. A cap then keeps at most 300 nodes per Biolink category, ranked by score, then coverage, then hop, so that populous categories (small molecules, genes) cannot crowd out rarer ones, and the category composition of the graph reflects the profile rather than the size of the categories in KG2c. Because the cap can remove the intermediate node through which another kept node was reached, a connectivity pass computes the nodes still reachable from the seeds through kept edges only, removes the others, and refills each category up to its cap with the next best candidates adjacent to the connected set; the pass is repeated at most three times and the result is flagged when it has not converged. The kept graph is written as GraphML with its node table (identifier, name, category, hop, coverage, score, degree, seeds and slides of origin) and edge table (predicate, weight, primary knowledge source, knowledge level), together with a summary that records the effective parameters, the number of nodes reached and kept per hop, the score floors, the connectivity passes and the structural metrics reported in the Results (components, diameter, seed-to-seed distances, vocabulary expansion).

### 2.5 Evaluation

#### 2.5.1 OCR task

The reading models were first compared on slide decks of the authors (subject-matter experts in muscular disorders) that combine microscopy images, scientific text and references, tables and flow charts. Four lightweight models that run below 16 GB of RAM without GPU (qwen2.5vl:3b, GLM-OCR, PaddleOCR and the general-purpose gemma4:e4b-it-qat) were tested on the FSHD1 deck, the most varied in image content. The OCR-specialised models transcribed text well but did not interpret biomedical figures, stylised text or annotations; the general-purpose models drew on their trained knowledge to name what the figures show, and qwen2.5vl:3b was the only one that interpreted the microscopy images, so it was retained. The prompt was then refined iteratively: a purely descriptive prompt extracted the content but conveyed too little of the scientists’ hypotheses, and was enriched with a generic biomedical persona that the user can focus on an area of interest. The model, the temperature and the profile were then compared in a controlled test (Supplementary Methods, *Reading-stage factor test*; Supplementary Tables S4 and S5).

#### 2.5.2 NER task

##### 2.5.2.1 Dataset Generation via stratified LLM-based augmentation

To construct the evaluation dataset, we employed a stratified sampling strategy based on the entity classes defined in the Biolink model ontology (Unni et al., 2022). The ontology was represented as a hierarchical structure (with mixins knowledge hence introducing horizontal linkage) from processing the Biolink YAML file. entity, serving as the top-level structural node and named thing representing the highest-level entity category were removed from entities of interest as uninformative structural roots. These two nodes were excluded from sampling because they function as ontology-level structural categories rather than target entity types. The objective was to create an evaluation set of biomedical entity mentions (queries), each paired with a target Biolink class. Every class of the Biolink model (release 4.4.4) except the two structural roots was a stratum with the same quota of ten mentions. Mentions were generated one class at a time by an open-weight model, gpt-oss-120b, and verified by a model of another family, deepseek-v4-flash; neither is the model evaluated in BiomedCAT. Both were accessed through an OpenAI-compatible endpoint (Albert, the French public-sector API). Each mention is one to four words long and was generated under the following constraints, each addressing a known bias of synthetic benchmarks:

- *Rare corpus representation bias*: every class is a stratum with the same quota, so rare classes count as much as frequent ones;
- *Model-family bias*: generator and verifier belong to different model families, and neither is the typing model under test (gemma);
- *Definition leakage*: the retrieval engine under test indexes each class’s definition and examples, so a mention must not repeat the class name, its examples, its aliases, or any word of five letters or more of its definition;
- *Canonical-form bias*: a surface-form quota per class covers four forms: canonical names, abbreviations or symbols, orthographic variants (plural, casing, hyphenation, common misspelling) and multi-word compounds;
- *Parent/child ambiguity*: every mention is re-classified by the verifier among the target class, its parent, its siblings and its children, and is kept only when the verifier returns the target; the verifier’s choice is stored so that near-misses can be analysed;
- *Cross-class duplicates*: global deduplication on the normalised string; surface variants of the same entity are allowed.

These rules rejected 895 generated mentions, and ten of the 157 classes kept none, the verifier assigning all their mentions to a child or sibling class (e.g. *disease or phenotypic feature* to *disease*) or to no class (bibliographic classes such as *journal article*). The resulting benchmark holds 1,356 verified mentions for 147 classes (412 canonical names, 382 abbreviations, 282 compounds and 280 variants; 144, 620, 340, 215 and 37 mentions at depths 2 to 6 of the class tree). It was then split, stratified by class and surface form, into a public slice of 1,075 mentions (147 classes), released with this article, and a private slice of 281 mentions (146 classes) that is never published, so that future models can be re-evaluated on mentions they cannot have seen in training. A canary string, a unique identifier embedded in every released file, allows a model that was trained on the public slice to be detected and redirected to the private slice. Every number reported in this article is computed on the public slice or on subsets of it. The generation, verification, split and canary scripts are in the companion repository (Data Availability Statement).

##### 2.5.2.2 Real gold set

SMEs created a list of terms (separately for each SME) from their slides that they expected the pipeline to recover. An expert bioinformatician and a computational scientist reviewed the terms and identified the best mapping given the scientific context of the slides. These term lists constitute our real gold standard, referred to as “gold”.

##### 2.5.2.3 Synthetic curated set

The labels of the public slice were audited by the bioinformatician expert against the Biolink class definitions, in two passes (18 and 22 September 2026), on 664 distinct mentions selected in the following order: the fixed 300-mention reference subset used for every configuration decision (Section 2.3.5); mentions on which three independent typers (Claude, DeepSeek and gemma-3-1b) agreed on another class than the label; one mention per class, to cover every class; and mentions added so that every reading scope (coarse stratum) reached the same number of scorable mentions. Model outputs were displayed only as a triage hint: the judgement bears on the label and definitions, not on the models. Each label received one of three verdicts: *ok*; *wrong*, with the class the expert would assign; or *ambiguous*, when several Biolink classes are defensible for the mention as written. Domain and range constraints were deliberately not used as a criterion, because the first curation pass found them missing or too narrow for several classes: the ranges of the Biolink model are inferred from a limited set of data sources that do not cover our use case. Generated mentions that name an identifiable person or a clinical case identifier were flagged as personal data, for which the abstention NONE is accepted as a correct answer in addition to the class. Classes without a definition in the model (such as *mammal* or *vertebrate*) were judged wrong automatically, because the expert could not verify the adequacy of a label to an absent definition. Plural mentions that do not denote the specific entity implied by the class definition were judged wrong and usually relabelled to the parent class. Ambiguous mentions received, when possible, every class they can take depending on context (each listed reading was checked manually; some acronyms remain too broad for their contexts to be listed). On the reference subset, 113 labels were confirmed, 125 were judged wrong and 62 ambiguous; over the 664 audited mentions, 298 were confirmed, 216 wrong (24 of them because the mention names no entity) and 150 ambiguous. Wrong labels are therefore not a marginal defect of LLM-generated benchmarks at this level of granularity. Absolute scores in this article are reported on the audited mentions, with the expert class as reference; the original labels are kept in the released file so that both readings can be reproduced.

##### 2.5.2.4 Metrics

Retrieval is scored by hit@k, the share of mentions whose expected class appears among the first *k* candidates of the ranked list, by the mean reciprocal rank and by the hierarchical distance between the first candidate and the expected class in the Biolink tree; typing is scored by accuracy, with 95% binomial confidence intervals, and two typing systems are compared on the same terms with an exact McNemar test on their discordant pairs. On the decks, reading is scored by the recall of the expert terms in the description, typing by accuracy on the recovered explicit terms, and linking by the share of recovered terms that receive the expert identifier (Supplementary Methods, *Expert gold of the slide decks*).

#### 2.5.3 Context-graph metrics

The context graphs are described by presence and structure metrics, not by a score of the hypothesis: the number of kept nodes and edges and of saturated categories, the vocabulary expansion (kept nodes per seed), the number of connected components, the share of seed pairs joined by a path inside the kept graph with their mean distance, the diameter, the Biolink category composition, the Jaccard overlap between the kept node sets of two graphs, and the recovery of a set of expected nodes written for each slide (pathway signatures) with their rank inside their Biolink category (Supplementary Methods, *Multi-scale case*).

## 3 Results

### 3.1 Overview of the workflow

BiomedCAT turns a slide deck into a user-defined contextual subgraph of RTX-KG2c in five stages (Figure 1). Each stage instantiates a distinct design pattern, and each pattern was chosen so that the workflow runs on a laptop with 16 GB of RAM: four small open models are served by Ollama and loaded one at a time, the previous model being released before the next one starts. (1) *Scientific description*: qwen2.5vl:3b reads every slide at temperature zero and writes a numbered list of dense scientific statements rather than a transcription, so that entities only shown in figures are named; the kinds of entities to name exhaustively come from the active profile. Each slide is screened by llama-guard3:1b before any other model reads it. (2) *Biolink categorization*: statements are split into sentences, candidate terms are extracted with a recall-first prompt, and every term is typed into one of the Biolink classes by gemma4:e4b-it-qat following a ReAct loop of at most three steps over three read-only tools (hybrid retrieval over class definitions and exemplars, class neighbourhood, exact lookup). Verdicts are constrained to the Biolink vocabulary and cached per document. (3) *Verification*: because a typing model can invent entities from their spelling, a term is kept only if a model of another family (llama3:8b) recognises it as a real biomedical entity and the Translator name resolver returns at least one candidate for it; the user may then review the list. (4) *Linking*: candidate identifiers are retrieved from the name resolver and the ARAX entity lookup, canonicalised to RTX-KG2c nodes through the local equivalence table and, for identifiers absent from it, the Node Normalizer without gene/protein or drug/chemical conflation, and gemma4:e4b-it-qat acts as a judge that selects one candidate or none. No identifier is ever generated by a model. (5) *Context graph*: the linked identifiers seed a two-hop expansion of the filtered RTX-KG2c executed inside DuckDB, where the profile sets the predicate, source and knowledge-level weights, the Biolink category scope and the edge directionality; pruning only bounds the graph for display and is not a ranking of hypotheses. Only entity names ever leave the machine, in stages (3) and (4), and both can be switched to a fully local mode: the resolver check of stage (3) is then dropped and stage (4) matches names and synonyms against the local KG2c table. The following subsections report each stage against its benchmark.

### 3.2 Choice of architecture

#### 3.2.1 Typing: empirical selection of the configuration

The typing configuration was selected through the twenty-one single-change experiments described in Methods (Section 2.3.5; Supplementary Table S6). Three findings shaped the final design. First, retrieval gained from evidence closer to how entities are written than to how Biolink defines them: a leg of exemplar mentions improved retriever hit@1 by 7 points and exact lexical matching improved hit@1 by 12 points on mentions from real slides, whereas a sparse BM25 leg, exemplars replaced by KG2c node names, and a longer candidate list (15 instead of 10) brought no measurable gain. Second, for a local model of about four billion parameters, more context did not mean better reasoning: the Biolink tree in the prompt, branch-level score aggregation, longer class definitions, a larger 12-billion-parameter model, additional reasoning passes and terse reasoning were neutral or detrimental. The only configuration above the threshold was a single call over a restricted menu (the ten best candidates plus the hierarchical neighbours of the top three), which reached 0.470 against 0.410 for the tool loop. The tool loop (ReAct) was nevertheless kept as the delivered configuration: once the labels were audited it scored higher than the single call (0.645 against 0.610 on the 664 audited terms, every expert reading of an ambiguous mention accepted), a difference that is not significant on paired terms (exact McNemar test, *p* = 0.08) and is reported as such (Section 2.5.2.3). Third, implementation details weighed as much as architecture: removing a chat role label from the prompt gained 5.7 points; sampling at the temperature recommended by the model vendor for gemma4:e4b-it-qat (1.0, top-k 64, top-p 0.95) gained 2.3 points on isolated terms but lost 25 to 31 points of exact recall on a real document, so greedy decoding was kept; and run-to-run reproducibility depended on how the model was served: with the model unloaded after each call on an otherwise idle machine, three runs of the tool loop on 50 benchmark terms gave identical verdicts, whereas keeping the model resident between calls lowered agreement to 82%, and complete runs on the 1,356 terms made with a resident model or while other experiments shared the machine agreed on 69 to 72% only; the published configuration therefore unloads the model after each call (two complete single-call runs under this setting agreed on 99.7%).

### 3.3 Evaluation

#### 3.3.1 Pipeline performance

Figure 2A gives the memory and time of each stage on one laptop (Intel i9-13900H, 64 GB of RAM, RTX 2000 Ada 8 GB). A one-slide deck takes 33 to 42 minutes end to end, the two-slide FSHD deck 59 minutes and the structure-only glycolysis image 16 minutes. Typing dominates (69 to 78% of the wall-clock time, 15 to 25 s per candidate mention); reading costs 108 to 114 s per slide, linking 3 to 6 s per mention and the context graph less than half a minute. Memory was sampled every second during the run of each of the eight documents (ranges over documents): the reading stage holds 3.3 GB of GPU memory (qwen2.5vl:3b) and 2.8 to 4.8 GB of system RAM above idle, typing 3.4 to 5.3 GB and 5.0 to 9.8 GB (gemma4:e4b-it-qat and, in turn, the llama3:8b check), linking 3.4 GB and 3.2 to 7.8 GB. In six of the eight runs the peak is not a model: the context-graph stage loads the RTX-KG2c edge table and reaches 6.0 to 9.6 GB of RAM for a few seconds, with the GPU idle; in the FSHD and Voie B runs the typing stage is higher. On a machine without GPU the model weights would move to RAM; adding them to the measured RAM gives an estimated peak of 10 to 13 GB, under the 16 GB design target, at the cost of CPU inference speed.

**Figure 2:**
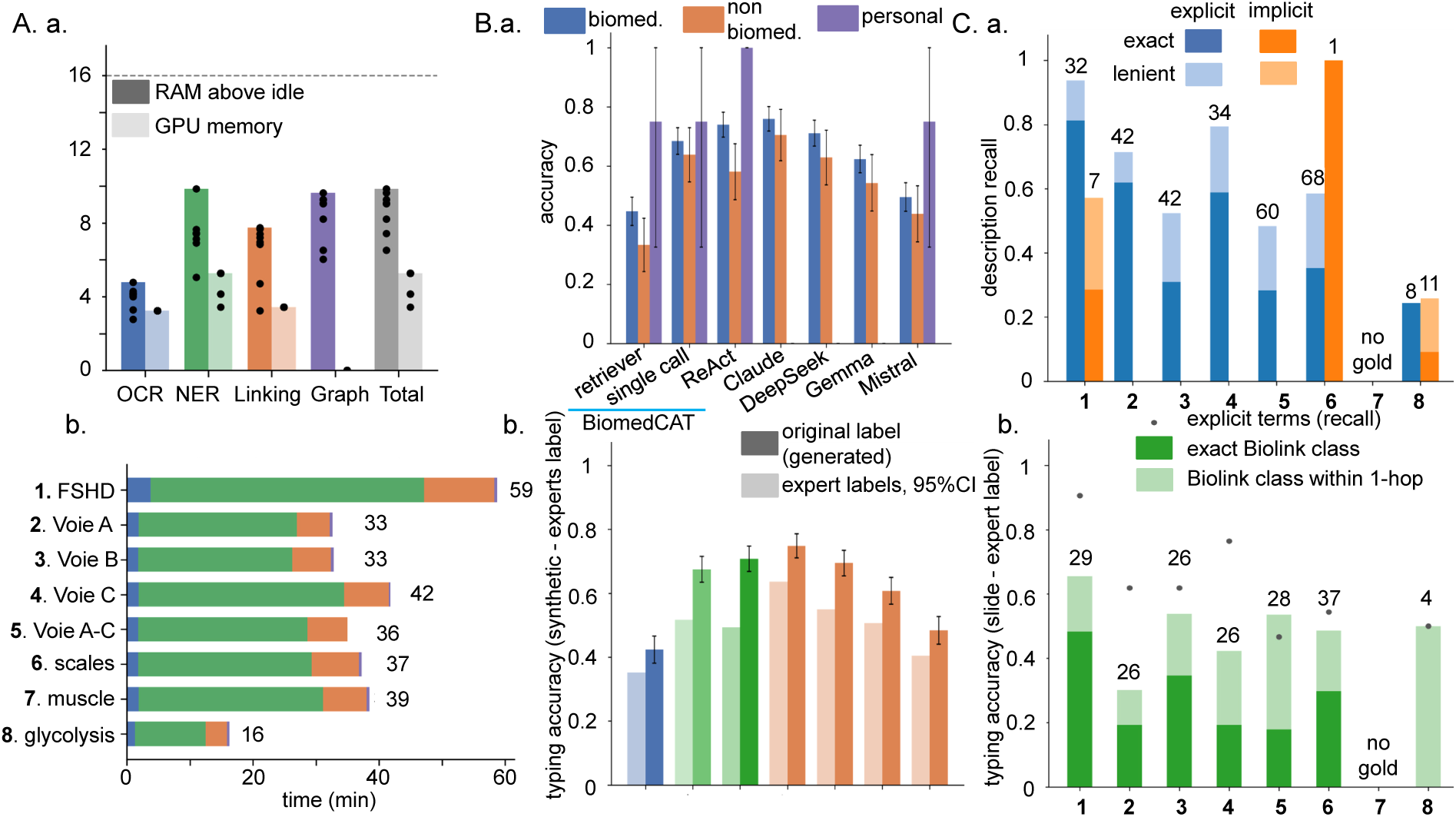
Resources, typing benchmark and expert-gold evaluation of the pipeline. Documents are numbered throughout: 1 FSHD (two slides, English), 2 Voie A, 3 Voie B, 4 Voie C, 5 Voies A–C, 6 FSHD1 multiscale summary slide, 7 muscle pathophysiology review table, 8 glycolysis structure image. **(A)** Resources of one run per document on one laptop (Intel i9-13900H, 64 GB of RAM, RTX 2000 Ada 8 GB). (a) Peak memory per stage: system RAM in use above the idle level (dark) and GPU memory in use (light); bars, maximum over the eight documents; dots, one document; dashed line, the 16 GB design target. (b) Wall-clock time per stage, total in minutes. **(B)** Biolink typing on the synthetic benchmark, 517 audited public mentions whose expert label is not ambiguous: BiomedCAT (retriever alone, top-1; single-call variant; ReAct agent, all on gemma4:e4b-it-qat, 4B parameters, local) against hosted models called one term at a time at temperature 0. (a) Accuracy against the expert labels by scope of the term: biomedical (n=412), non-biomedical (n=105; information, administrative, attribute, activity, physical and planetary classes) and personal data (n=4, named people, a subset of the biomedical terms, for which NONE is also accepted); error bars, 95% binomial interval. (b) Accuracy against the original LLM-generated labels (light) and against the expert labels (dark, 95% binomial interval), any listed expert class accepted and NONE expected when the mention is not an entity. **(C)** Evaluation against the harmonized expert gold of the documents (Supplementary Methods, *Expert gold of the slide decks*; document 7 has no expert gold). (a) Recall of the expert terms in the text produced by the reading stage, explicit terms (written on the slide, blue) and implicit terms (expected by the expert, not written, orange), under exact string matching (dark) and lenient matching that accepts French/English cognates (light); numbers, terms per document. (b) Typing accuracy on the explicit terms recovered by the extraction (numbers): exact expert Biolink class (dark) and class at hierarchical distance at most one (light); dots, share of explicit terms recovered by the extraction. Source data: data/publication/figures_data/.

#### 3.3.2 Pipeline quality against the expert gold

Each stage is scored against the harmonized expert gold of the documents (Figure 2C) and, for typing, against the audited synthetic benchmark (Figure 2B).

##### 3.3.2.1 Reading: recall of the expert terms

Against the harmonized expert gold of the seven decks (Supplementary Methods, *Expert gold of the slide decks*), the reading stage reproduces 81% of the explicit terms of the English FSHD deck under exact string matching and 94% under the lenient matching that accepts cognates, and 57% of the seven implicit terms the expert expected without their being written on the slide (Figure 2C.a; Supplementary Table S1). On the French decks the exact recall falls to 28 to 62%, and 48 to 79% with lenient matching: the description is written in English, and French terms without an English cognate (*puissance*, *récupération*, *noyau*) are counted as missed even when their concept is present. The structure-only glycolysis image is the failure case: the vision model described a nucleotide-synthesis pathway instead of the drawn glycolytic steps, and only 2 of 8 explicit and 3 of 11 implicit terms are recovered. Reading is deterministic at temperature zero: the two FSHD runs give identical recall.

##### 3.3.2.2 Typing on the decks

On the explicit expert terms that the extraction recovered (26 to 37 per deck), the typing agent assigns the exact expert Biolink class in 48% of cases on the FSHD deck and 18 to 35% on the French decks; a class at one step in the Biolink tree (parent or child of the expert class) is reached in 66% and 31 to 54% respectively (Figure 2C.b). The extraction itself recovers 91% of the explicit FSHD terms and 47 to 77% on the French decks, where the agent must type an English paraphrase of a French label. The errors are of two kinds, mechanism rather than entity (a disease typed as a pathological process) and granularity (nucleic acid entity against regulatory region), which the one-step criterion separates from outright misses.

##### 3.3.2.3 Typing on the synthetic benchmark

The typing stage was evaluated on the public slice of the synthetic benchmark (1,075 mentions, 147 classes; 340 canonical names, 314 abbreviations, 211 orthographic variants and 210 compounds; 114, 492, 271, 169 and 29 mentions at depths 2 to 6 of the class tree), on its 300-mention reference subset for the configuration decisions, and on the 664 audited mentions for the absolute scores (Methods, Section 2.5.2.3). On the public slice, the hybrid retriever alone ranks the expert class first for 0.333 of the mentions (hit@5 0.576, hit@10 0.655, mean reciprocal rank 0.447, mean hierarchical distance of the first candidate to the labelled class 2.8), and the ReAct agent built on it reaches 0.44 (*±*0.03) against the original labels, the single-call variant 0.48. Figure 2B compares the typing stage with hosted models on the 517 audited public mentions whose expert label is not ambiguous (of the 150 mentions judged ambiguous, only the three that name a person are kept, NONE being accepted for them). Against the original LLM-generated labels, BiomedCAT’s ReAct agent scores 0.49 and the single-call variant 0.52; against the expert labels, 0.71 (95% CI *±*0.04) and 0.68. The hosted models follow the same shift: Claude Opus 5 from 0.64 to 0.75, DeepSeek-V4-Flash from 0.55 to 0.69, gemma-4-31b from 0.51 to 0.61, mistral-small-24b from 0.40 to 0.48. The retriever alone reaches 0.42. On paired terms (exact McNemar test), the 4-billion-parameter local agent does not differ significantly from Claude Opus 5 (*p* = 0.10) or DeepSeek-V4-Flash (*p* = 0.63) and is above the two open-weight models of 24 and 31 billion parameters (*p <* 0.001). By scope, the agent scores 0.74 on the 412 biomedical terms and 0.58 on the 105 non-biomedical terms (information, administrative and attribute classes); on the four personal-data terms it answers NONE four times, which the privacy rule requires, where none of the hosted models does so. Including the ambiguous terms with any listed reading accepted, the agent scores 0.65 on all 664 audited mentions, Claude Opus 5 0.69 and DeepSeek-V4-Flash 0.63 (Supplementary Table S3); on this larger set the gap to Claude Opus 5 is significant (*p* = 0.047).

##### 3.3.2.4 Linking: online services against a fully local mode

On the FSHD deck, with reading and typing identical in both modes, the online linking (Name Resolver and ARAX candidates selected by the local judge) assigns the expert identifier to 65% of the 23 found terms that have one in RTX-KG2c, against 26% for the fully local mode, which matches names against the local KG2c table only (Supplementary Figure S1 and Supplementary Methods, *Linking: online services against a fully local mode*). Linking is the only stage where leaving the machine buys accuracy; the user chooses it per run.

### 3.4 Study case: gold standard slides in FSHD

The FSHD1 inflammation deck of the collaborating team contains three slides that each draw one pathway from DUX4 to muscle vulnerability, at a different scale: slide A, molecular (aberrant transcripts and retroelements sensed as double-stranded RNA by RIG-I, MDA5 and TLR3, interferon and ISG induction, MHC-I presentation); slide B, organelle (mitochondrial dysfunction, reactive oxygen species, lipid oxidation and mtDNA damage, HMGB1 and inflammasome signalling, bioenergetic deficit and fatigue); slide C, tissue (damaged fibres, macrophages and fibro-adipogenic progenitors, fibrosis and fat replacement, force transmission, MRI). A fourth slide draws the three pathways together, and the deck closes on a multi-scale summary slide. The five slides were processed by the BiomedCAT pipeline and their linked entities seeded a context graph under the biochemical profile (2 hops, per-category cap 300). We asked three questions of these graphs: does the expansion follow the biological scale presented in the slide, how does the structure of the kept graph change with the breadth of the hypothesis, and does the graph connect DUX4 to the downstream events the researchers drew?

#### 3.4.1 Context graphs of the five slides

##### 3.4.1.1 Recovery of the pathway signatures

For each pathway we listed ten RTX-KG2c nodes that a reader of the slide expects in its neighbourhood (Reactome pathways and reactions, GO processes, genes; Supplementary Table S7) and that are seeds of no deck, so that every occurrence is produced by the expansion. The recovery matrix is diagonal (Figure 3c, Supplementary Table S9). The slide-A graph holds nine of the ten antiviral-sensing nodes (*DDX58/IFIH1-mediated induction of interferon-α/β* at rank 8 of 300 kept pathways, *TRAF6 mediated IRF7 activation* at rank 11, *ISG15 antiviral mechanism* at 18, the *TLR3 cascade* at 49; median rank percentile 0.16), one mitochondrial node and three remodelling nodes. The slide-B graph holds none of the antiviral nodes and five of the mitochondrial ones, all at hop 1 and prominent (*4-hydroxynonenal* rank 1 of the small molecules, *reactive oxygen species metabolic process* rank 5, *mitochondrial complex I* rank 8; median percentile 0.06). The slide-C graph holds eight of the ten remodelling nodes (*TNF-α*, *tumor necrosis factors*, *chemokine CCL2* and the *execution phase of apoptosis* within the first five of their categories; *extracellular matrix organization* and *degradation of the extracellular matrix* at ranks 35 and 34 of the pathways; median percentile 0.07). The off-diagonal cells are not empty, but their hits sit low in their categories (median percentiles 0.35 to 0.58). Two of the slide-B signature nodes, the HMGB1 and NLRP3 genes, are written on slide B but were not linked by the pipeline; the expansion brings them back at hop 1.

**Figure 3:**
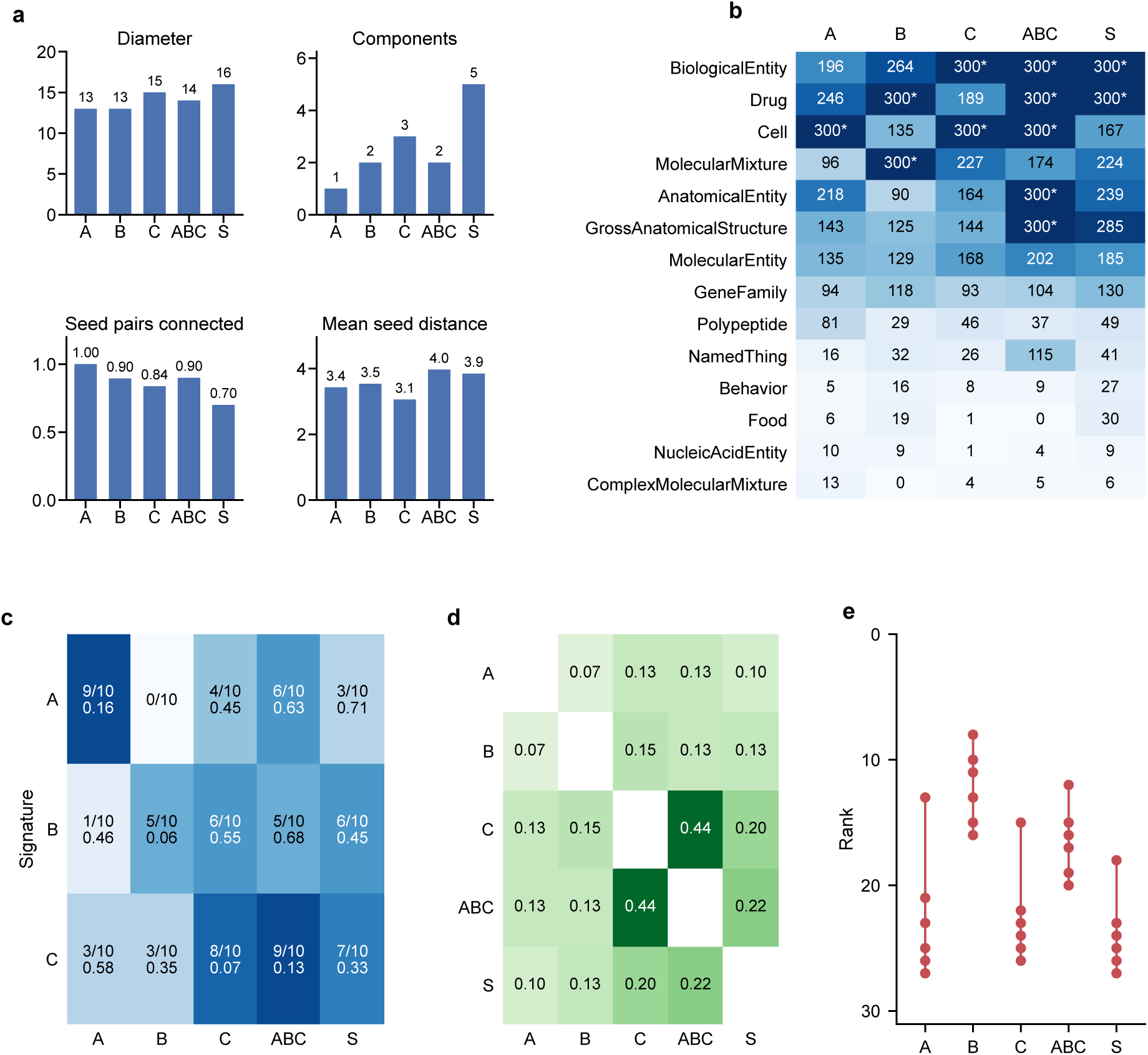
Context graphs of the three pathway slides, of their integration and of the summary slide of the FSHD1 inflammation deck. All graphs were extracted with the biochemical profile (2 hops, per-category cap of 300 nodes). Slides: A, molecular scale (innate antiviral sensing); B, organelle scale (mitochondria and redox); C, tissue scale (immune infiltrate and fibro-adipogenic remodelling); ABC, the slide that draws the three pathways together; S, the multi-scale summary slide. (a) Structure of the kept graph: diameter (hops), number of connected components, share of seed pairs joined by a path inside the kept graph, and mean shortest path between connected seed pairs (hops). (b) Kept nodes per Biolink category, for the categories in which the five graphs differ; the thirteen most populated categories (genes, proteins, transcripts, small molecules, pathways, processes, diseases, phenotypes, …) are at the cap of 300 in every graph and are not shown; an asterisk marks a category at the cap. (c) Recovery of the pathway signatures A, B and C: for each pathway, ten RTX-KG2c nodes that a reader of the slide expects (Reactome pathways and reactions, GO processes, genes; Supplementary Table S7) and that are seeds of no slide. In each cell, the first line is the number of signature nodes present in the kept graph, the second the median, over those present, of the rank percentile of the node inside its own Biolink category (0 = first of its category, 1 = last); colour, share of the ten nodes present. (d) Jaccard index of the kept node sets of two graphs (nodes common to both over nodes of either); the ABC graph shares 67% of its nodes with the union of the A, B and C graphs. (e) Rank, among the 300 kept MolecularActivity nodes of each graph (1 = best score), of the six Reactome reactions of DUX4 target activation (*Expression of DUX4 in the zygote*; *DUX4 binds* KDM4E, LEUTX, HERVL LTR, MaLR, HSATII), none of which is a seed; all six are present in every graph; dots are the3s4ix ranks and the bar their range.

##### 3.4.1.2 Structure of the kept graphs

The three single-pathway slides give graphs of the same size (5,491 to 5,604 nodes, 51,748 to 53,136 edges: the per-category cap saturates 14 or 15 of the 31–32 categories in every case), so size does not separate them (Supplementary Table S8). The categories below the cap follow the slide (Figure 3b): cells fill their 300 places in the graphs of slides A and C but only 135 in that of slide B, molecular mixtures (mitochondrial complexes, lipid preparations) reach the cap only for slide B, and gross anatomical structures only for the integrated slide. Connectivity, in contrast, separates them (Figure 3a). The molecular slide A gives one connected component in which every pair of its 20 seeds is connected (190/190, mean seed-to-seed distance 3.4 hops, diameter 13); the organelle slide B two components (153/171 pairs, diameter 13); the tissue slide C three components (231/276 pairs, diameter 15), although its seeds are the closest to one another on average (3.1 hops). The slide that integrates the three pathways gives the largest graph (6,080 nodes, 74,541 edges, 18 saturated categories, vocabulary expansion 304 nodes per seed against 234–289), with the largest mean seed-to-seed distance (4.0 hops) at a diameter of 14. The summary slide, whose 25 seeds span 17 Biolink types from the nucleus to the clinic, gives the most fragmented graph (5 components, 210/300 seed pairs connected, diameter 16). From slide A to the summary slide the diameter grows from 13 to 16 and the number of components from 1 to 5, while the share of nodes at hop 1 stays between 0.27 and 0.40. The kept node sets of the three single pathways overlap little (Jaccard 0.07 to 0.15; Figure 3d); the graph of the integrated slide is closest to that of slide C (Jaccard 0.44), and 67% of its nodes are present in at least one of the A, B and C graphs, the remaining third being nodes that none of the single-pathway graphs reached.

##### 3.4.1.3 Paths from DUX4

The six Reactome reactions of DUX4 target activation (*Expression of DUX4 in the zygote*, *DUX4 binds* KDM4E, LEUTX, HERVL LTR, MaLR, HSATII), which are seeds of no slide, are present in all five graphs, within the first 27 of the 300 kept molecular activities, and within the first 16 for slide B (Figure 3e). Figure 4a–c shows a window on each single-pathway graph, and Figure 4d reads, for each pathway, the best-weighted path from DUX4 to the pathway signature (the stage’s own score: product of profile weights divided by the degree penalty of intermediate nodes). In the graph of slide B, DUX4 reaches HMGB1 in one step through a *physically interacts with* assertion from IntAct; the same edge is the path in the slide-C graph. In the integrated slide, DUX4 reaches the antiviral innate immune response through RELA (NF-*κ*B p65), NLRP3 through a shared cytosolic localisation, and the execution phase of apoptosis through a cell-differentiation process; the summary slide reaches the *DDX58/IFIH1* pathway through APP and the collagen-containing extracellular matrix through TGM2. Inside each kept graph the three signatures lie two to four hops from one another, except in the slide-B graph, where the antiviral signature is absent altogether (Supplementary Table S10).

**Figure 4:**
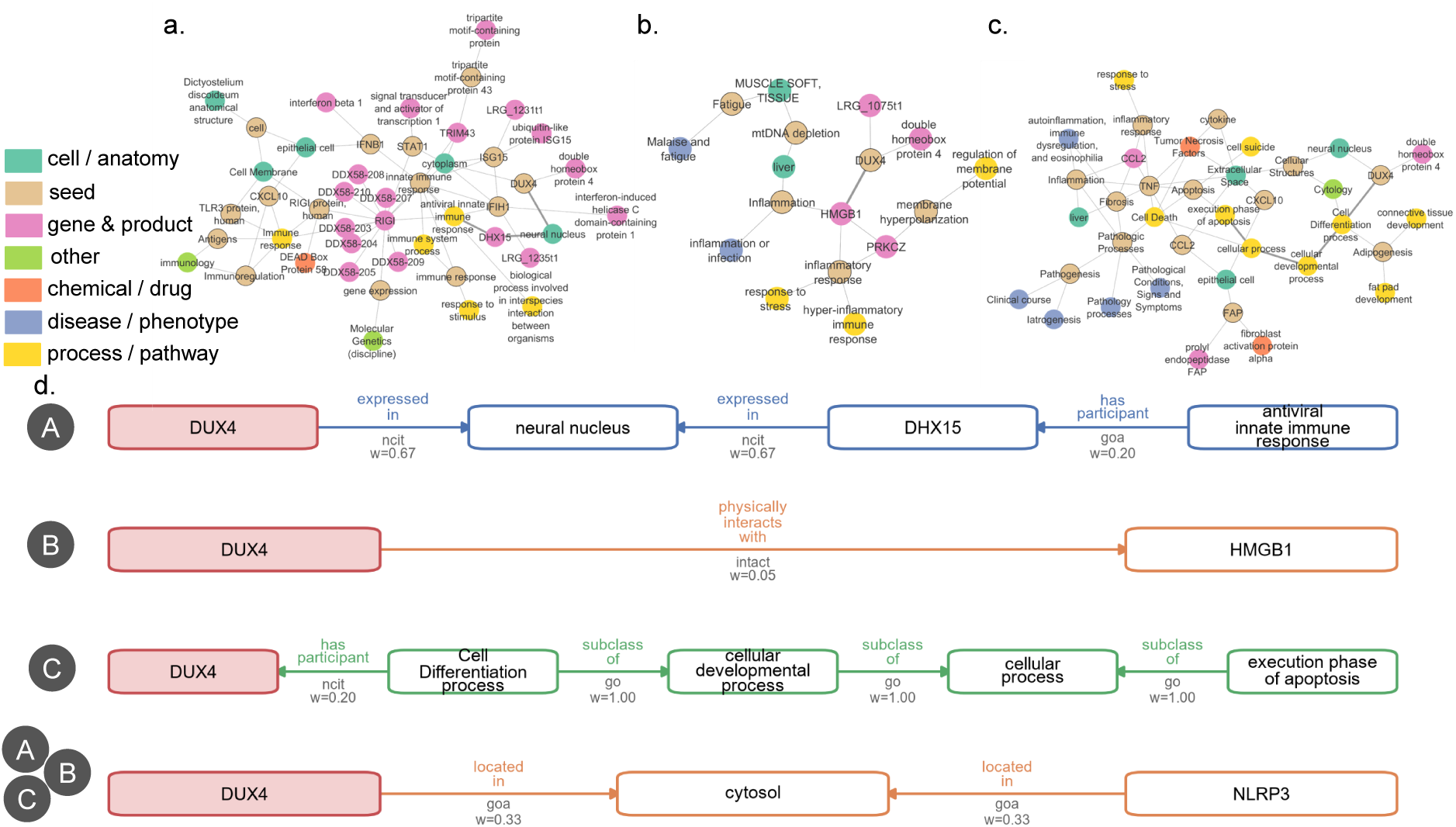
From DUX4 to each pathway. (a–c) A window on the context graph of each single-pathway slide of the FSHD1 inflammation deck (a, slide A, molecular scale; b, slide B, organelle scale; c, slide C, tissue scale; biochemical profile). The window holds the seeds of the slide, the two best-scored kept neighbours of every seed, the ten best-scored nodes of the graph and the nodes of the best-weighted path from DUX4 to the slide’s own pathway signature, with the edges of the kept graph between these nodes; for legibility, only the largest connected component of each window is drawn. Shown: 46, 17 and 38 nodes (88, 17 and 58 edges) of the 5,491, 5,500 and 5,604 kept nodes, including 15 of the 20, 6 of the 19 and 14 of the 24 seeds; the seeds left out include *double-stranded RNA* for slide A, *mitochondrion*, *4-hydroxynon-2-enal*, *lipid oxidation* and *reactive oxygen species biosynthetic process* for slide B, and *macrophage*, *fibro/adipogenic progenitor cell*, *tissue remodeling* and *necrosis* for slide C (full windows: Supplementary Methods, *Best-weighted paths from DUX4*). Node colour: seed (beige), gene or gene product (pink), biological process or pathway (yellow), chemical or drug (orange), cell or anatomical entity (green), disease or phenotype (blue), other Biolink classes (light green); darker edges, the path from DUX4. (d) Best-weighted path from DUX4 to a pathway signature, read as a chain of RTX-KG2c assertions (predicate above each edge, primary knowledge source and profile weight below; arrows in the direction of the assertion): A, in the slide-A graph to the antiviral signature; B, in the slide-B graph to the mitochondrial signature (the same single edge is the path in the slide-C graph); C, in the slide-C graph to the remodelling signature (the same path in the graph of the integrated slide); A+B+C, in the graph of the slide that integrates the three pathways, to the mitochondrial signature. The path cost is the stage’s own score (product of profile edge weights divided by the degree penalty of intermediate nodes). The path to HMGB1 is a single assertion of the IntAct interaction database; the paths through *neural nucleus* and *cytosol* are co-localisation edges from the Gene Ontology Annotation database (GOA) and NCIT (see Discussion). The other paths are listed in the Supplementary Methods.

#### 3.4.2 Compounds tested in FSHD in the graphs of the FSHD deck

To relate the context graphs to drug discovery, we looked for 21 compounds that have been tested in FSHD, from investigational agents to repurposed marketed drugs and supplements (Supplementary Methods, *Compounds tested in FSHD*), in the graphs of the two-slide FSHD deck built under the six profiles (Table 1). The deck names the genetic and epigenetic mechanism (D4Z4, DUX4, SMCHD1, DNMT3B) and the disease but no treatment, so every compound found is brought by the expansion. Seven compounds have no node in RTX-KG2c, among them the three agents that target DUX4 (two siRNA conjugates and an epigenetic editor): no expansion of this knowledge graph can return them, which bounds what a graph-based approach can propose for a disease whose therapeutics are recent. Of the 14 others, 12 are reached within two hops in at least one graph, but few are kept: two to four per profile, and none under the genetic-determinants profile, whose scope excludes chemical entities. Prednisone and testosterone, repurposed for their anti-inflammatory and anabolic effects, are kept at hop 1 under four profiles, while creatine and the antioxidant supplement depend on the profile. Losmapimod, the p38*α/β* MAPK inhibitor that reached phase 3 in FSHD, is reached at two hops in five graphs but kept only in the uniform one, and its target MAPK14 is reached but pruned in every graph (rank 680 to 958 among the 16,000 to 18,500 genes reached, of which 300 are kept). The graphs thus place compounds tested in FSHD within two hops of the deck’s entities, but rank them by their connection to what the slides say, not by therapeutic relevance; a drug-discovery reading needs a seed or a profile that points to the target.

**Table 1:**
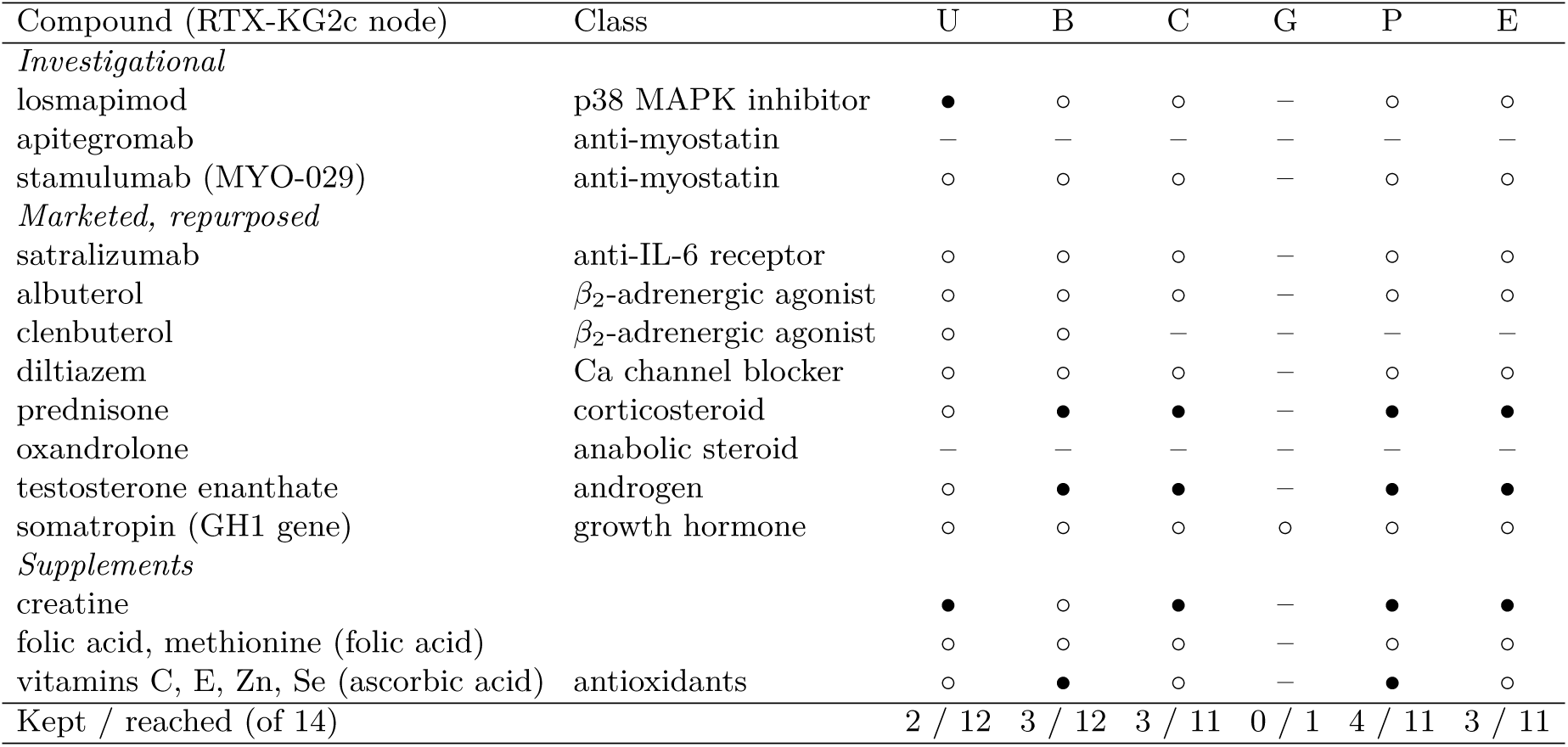
Compounds tested in FSHD in the context graphs of the FSHD deck under the six profiles. *•*, kept in the graph; *◦*, reached within two hops but pruned (retention floor or category cap); –, not reached. U, uniform; B, biochemical actions; C, clinical mechanisms; G, genetic determinants (its scope excludes chemical entities); P, pharmacological intervention; E, epidemiological risk. Compounds tested in combination are matched on one component, and somatropin on the GH1 gene, its only node in RTX-KG2c (node in brackets). Seven compounds of the list have no node in RTX-KG2c: delpacibart braxlosiran, ARO-DUX4 and EPI-321, which target DUX4, and GYM329, ACE-083, ATYR1940 and SFL-0821. Source data: data/eval/fshd_presence_pub.json.

| Compound (RTX-KG2c node) | Class | U | B | C | G | P | E |
|---|---|---|---|---|---|---|---|
| *Investigational* |  |  |  |  |  |  |  |
| losmapimod | p38 MAPK inhibitor | ● | ○ | ○ | – | ○ | ○ |
| apitegromab | anti-myostatin | – | – | – | – | – | – |
| stamulumab (MYO-029) | anti-myostatin | ○ | ○ | ○ | – | ○ | ○ |
| *Marketed, repurposed* |  |  |  |  |  |  |  |
| satralizumab | anti-IL-6 receptor | ○ | ○ | ○ | – | ○ | ○ |
| albuterol | *β*<sub>2</sub>-adrenergic agonist | ○ | ○ | ○ | – | ○ | ○ |
| clenbuterol | *β*<sub>2</sub>-adrenergic agonist | ○ | ○ | – | – | – | – |
| diltiazem | Ca channel blocker | ○ | ○ | ○ | – | ○ | ○ |
| prednisone | corticosteroid | ○ | ● | ● | – | ● | ● |
| oxandrolone | anabolic steroid | – | – | – | – | – | – |
| testosterone enanthate | androgen | ○ | ● | ● | – | ● | ● |
| somatropin (GH1 gene) | growth hormone | ○ | ○ | ○ | ○ | ○ | ○ |
| *Supplements* |  |  |  |  |  |  |  |
| creatine |  | ● | ○ | ● | – | ● | ● |
| folic acid, methionine (folic acid) |  | ○ | ○ | ○ | – | ○ | ○ |
| vitamins C, E, Zn, Se (ascorbic acid) | antioxidants | ○ | ● | ○ | – | ● | ○ |
| Kept / reached (of 14) |  | 2 / 12 | 3 / 12 | 3 / 11 | 0 / 1 | 4 / 11 | 3 / 11 |

## 4 Discussion

Existing KG-grounding methods retrieve information from curated databases in response to text prompts. BiomedCAT starts instead from the researcher’s own material: it reads slides with a small vision-language model, types and links the entities they name with local models and deterministic services, and returns a subgraph of RTX-KG2c shaped by the researcher’s profile. The researcher thus explores the biological neighbourhood of their own hypothesis on their own machine, with every seed traceable to the slide that introduced it and every edge to its primary source.

### 4.1 Choice of lightweight models for local usage

Every component was chosen to run on a laptop without a GPU budget: four open models of one to eight billion parameters served by Ollama and loaded one at a time, prompts kept short (the design log shows that longer context, the Biolink tree in the prompt or longer definitions, made the 4-billion-parameter typing model worse, not better; Supplementary Table S6), and a retrieval index of a few tens of megabytes in place of an in-memory copy of the field. Under these constraints the typing stage reaches 0.71 against the expert labels, 4 points below Claude Opus 5 (0.75) and above DeepSeek-V4-Flash (0.69), neither difference being significant on paired terms (Figure 2B.b). The whole run of a one-slide deck takes 33 to 42 minutes and peaks at 9.8 GB of system RAM above idle, with at most 5.3 GB of GPU memory (Figure 2A).

Running locally also makes the energy budget of a run easy to bound. With the laptop drawing between 60 W (CPU, idle GPU) and 200 W (GPU at full load), a one-slide deck costs 0.03 to 0.14 kWh and the two-slide FSHD deck 0.06 to 0.20 kWh, that is less than 4 g of CO_2_e at the 2025 carbon intensity of French electricity and 6 to 35 g at the European average (Supplementary Methods, *Resource measurement*).

Reproducibility was treated as a parameter to select, not a property to assume. All models run with greedy decoding and a fixed seed and are released from memory after each call, and every run records the model identifiers, the Biolink release, the profile and the versions of the remote services (run record of each deck and environment.json; Data Availability Statement). Under these settings the reading stage is deterministic (the two FSHD runs give identical recall), and two complete runs of the document pipeline on the FSHD deck gave identical typed and linked entities, and the publication runs, four weeks later, reproduced the same 175 typed mentions. The tool loop is the element most sensitive to execution conditions: with the model unloaded after each call on an idle machine, three runs on 50 benchmark terms gave identical verdicts, but keeping the model resident lowered agreement to 82%, and complete benchmark runs made with a resident model or while other experiments shared the machine agreed on 69 to 72% (Supplementary Table S6, E20). The context graph is reproducible up to ties: two builds differ only in which nodes of equal score fill the last places of a saturated category (Methods, Section 2.4).

#### 4.1.1 Choice of the agentic architecture

The selection experiments (Results, *Choice of architecture*) also say what a four-billion-parameter model can use. Every change that gave it more to read (the Biolink tree, longer definitions, more candidates, more reasoning passes) was neutral or harmful, and the two changes that helped brought the evidence closer to how entities are written (exemplar mentions, exact lexical matching) or removed noise from the prompt (the chat role label). The tool loop was kept over the single call although it scored lower against the original labels, because the audited labels reversed the order and because its three tools are what lets a verdict be traced to a definition, a lookup or a neighbourhood; its reproducibility, in turn, requires the published serving setting (model unloaded after each call) and a machine not shared with other model jobs.

The three-stage design separates recall from precision. The reading stage is tuned for recall (a description of every slide rather than a transcription, so that entities only drawn in figures are named) and the extraction prompt keeps every candidate; the typing agent and the existence gate are tuned for precision, because in biomedical use a false positive costs more than a miss: an invented gene that enters the context graph seeds a neighbourhood that does not exist. On the FSHD deck the extraction recovers 91% of the explicit expert terms at a mention precision of 0.53 (0.60 on distinct mentions); the gate rejects 32 of the 34 invented terms of the adversarial suite at the cost of two real terms, *repeats* and *upper-arms* (union mode; Supplementary Table S2), and 9 of the 34 invented terms that reach the typing agent in isolation, gate off, still receive a class, none an identifier. The agent’s abstention (NONE) is the other half of this precision: it is the verdict the audit expects for strings that name nothing, and the one the hosted models rarely give.

### 4.2 Images as input modality for hypothesis driven data

The input format was chosen to minimise the researcher’s effort: the pipeline starts from what scientists already produce, their slides and figures, exported as PDF or images, with no annotation, no template and no query language to learn. While existing systems are designed for clean plain-text sources such as PubMed abstracts, as in PubMed Knowledge Graph 2.0, researchers communicate their findings and hypotheses through presentation slides, which most of the time combine text blocks, figures, tables, and charts. Layout matters all the more in biomedical slides, whose gene symbols, chemical subscripts and domain-specific abbreviations are short tokens: the corruption of a single character is enough to change the entity, as the reading-stage factor test shows (*D4Z* for D4Z4, *FSD1* for FSHD1; Supplementary Methods). The workflow is designed for the few slides that state a working hypothesis, not for the full content of a publication; its running time grows with the number of mentions (Section 4.7).

### 4.3 From explicit knowledge to a user-defined mental model

Publications and patents record the outcome of research, often without the intermediate reasoning and experimental context, which are left out for publication or for the protection of intellectual property. Internal presentations keep part of this context: the pathways a team is considering, its preliminary observations and the links it has not yet tested, a form of tacit knowledge (Mitchell et al., 2022; Li et al., 2026). BiomedCAT takes these documents as its input and returns the part of the knowledge graph that they point to, so that a researcher’s working model of a mechanism can be set against what the graph already holds, and the gaps between the two can be seen.

### 4.4 Memory performance and data privacy

Because the models are loaded one at a time, each released before the next starts, the peak memory of a run is set by its largest single component rather than by their sum: the context-graph stage, which loads the RTX-KG2c edge table, in six of the eight measured runs, and the typing stage in the two others (Figure 2A). Without a GPU the model weights move to RAM, for an estimated peak of 10 to 13 GB, under the 16 GB target at the cost of inference speed; the model server (Ollama) selects the device itself, and the pipeline has a single code path.

Local execution is a requirement of the use case, not a convenience. Research presentations hold unpublished hypotheses, preliminary data and, at times, patient-level information; a tool that sends them to a hosted model is not usable for grant preparation or for experimental reasoning before publication. In BiomedCAT the slides, their descriptions and the typed mentions never leave the machine. Only bare entity names are sent out, at two points (the existence check and the candidate lookup of the linking stage), and both can be switched off: the fully local mode keeps reading and typing unchanged and links by exact name against the local KG2c table. The price of that mode is coverage: the online resolvers index every identifier and synonym of the Translator ecosystem, and linking accuracy on the FSHD deck falls from 0.65 to 0.26 without them (Supplementary Figure S1). Terms typed into person-denoting classes, and strings that look like identifiers or contact data, are never sent out in either mode. The user thus chooses, per run, between full privacy and the coverage of international standards, with the same reading and typing in both cases.

### 4.5 Evaluation

Evaluating a pipeline that reads, types and links biomedical entities requires a benchmark at the granularity of the target model, here the classes of the Biolink model.

#### 4.5.1 Evaluating biomedical entity typing

In the absence of a manually annotated corpus of mentions typed against the Biolink model, we built the typing benchmark with language models, under constraints that respect the granularity of the Biolink model (Methods). Sampling all depths of the class tree (2 to 6), mixins included, makes the evaluation cover both the assignment of a generic mention to a broad class and the discrimination between specific subclasses.

The expert audit qualifies this design. On the 300-mention reference subset, only 113 labels were confirmed, 125 were judged wrong and 62 ambiguous, although every mention had passed cross-family verification. Inspection of the verdicts points to two mechanisms. The generator often writes a mention for what the class is about rather than for an instance of it, which is a label error. And for a short mention taken out of context, several Biolink classes are frequently defensible (a drug name is also a small molecule; a cohort study is also a study), which is a property of the task that the benchmark can expose but not remove.

Three consequences follow for reading our results. Differences between configurations measured against the original labels remain informative, because the label error is the same for every configuration, but absolute scores are only meaningful on the audited mentions. Synthetic mentions are short strings without a sentence, whereas the pipeline types mentions read from a document in context; the gain of the exact lexical leg (+0.5 point on synthetic mentions, +12 points on the mentions of a real deck) shows how far the two settings can diverge. Finally, a benchmark released publicly will enter future training corpora; the private slice and the canary string are our provision for re-evaluating future models on mentions they have not seen.

Future work will add a second annotator to the audit, to measure agreement on the ambiguous cases, and will pair the synthetic mentions with mentions drawn from the knowledge graph and from real documents.

#### 4.5.2 Linking

We did not ask the generative model to produce identifiers. An identifier such as NCBI-Gene:100288687 carries no meaning that a language model could have learnt from text: its digits are not associated with the concept they index, and asking a model for one invites a well-formed but invented answer that no downstream check can distinguish from a correct one. The linking stage therefore reverses the roles: deterministic services and tables propose candidates (the Name Resolver, ARAX, the local KG2c name table), each shown with its label, its Biolink category and its presence in the graph, and the model only chooses among them or abstains. Its contribution is the disambiguation that a lexical lookup cannot make, choosing the human gene among orthologues or the process among the entities that share its name, in the light of the sentence and of the class decided at typing.

Linking is better online because coverage is a property of the index, not of the model: the Name Resolver indexes about 330 million concepts with their names and synonyms (Babel release of July 2026), against the names and synonyms of the filtered KG2c node table locally, and the difference is visible on the FSHD deck (Results, *Linking*). Six of the 29 found expert terms have no identifier in either KG2c or Node Normalizer; they are not a failure of the linker but a limit of the current vocabularies, and each Babel release extends them. The versions of the services are recorded with every run so that a later run can be compared on the same footing.

### 4.6 Study case: a graph per hypothesis

#### 4.6.0.1 A graph per hypothesis, not a graph per disease

The three pathway slides share one gene, one disease and the word inflammation, and a disease-centred expansion of RTX-KG2c would have given them the same neighbourhood. Seeding the graph with what the slide actually says gives three graphs that overlap by 7 to 15% of their nodes and that each recover their own pathway’s expected nodes near the top of their categories while leaving the other two pathways’ nodes near the bottom (Figure 3c, d). The context graph therefore encodes the focus of the hypothesis and not only its subject, which is what makes it usable as a mental-model companion: a reader of slide B will find mitochondrial complex I, 4-hydroxynonenal and the reactive oxygen species metabolic process in the first ranks and will not have to filter out interferon signalling, and conversely.

#### 4.6.0.2 What the structure of the kept graph says about the slide

Because the per-category cap bounds every graph to about the same size, size is not the measure of a slide’s breadth; connectivity is. A slide that stays at one scale gives one component in which every seed reaches every other (slide A); a slide that spans scales gives seeds that do not reach one another within two hops of RTX-KG2c, hence several components and a larger diameter: the cell- and tissue-level seeds of slide C (macrophage, fibro-adipogenic progenitor, mechanical stress, tissue regeneration) are densely connected to each other but not to all of its molecular seeds, and the summary slide spans the most levels. This is a property of the knowledge graph as much as of the slide: the organism-level and tissue-level vocabularies of RTX-KG2c (anatomy, phenotypes, procedures) and its molecular vocabularies (genes, reactions, small molecules) are connected mostly through disease nodes, so a hypothesis that crosses scales crosses a sparse region of the graph. We report these numbers as presence metrics and do not read them as a score of the hypothesis: a fragmented graph tells the user where the knowledge graph does not yet bridge the levels the slide bridges, which is itself the kind of gap a mechanistic hypothesis is written to fill.

#### 4.6.0.3 Reading a composite slide

The integrated slide, which repeats the three pathways in a denser layout, was the hardest to read: 39 of its 64 mentions were linked against 55 to 77 for the single slides, and its graph is closer to slide C’s (Jaccard 0.44) than to A’s or B’s, because the tissue-level labels of the composite figure (macrophage, fibrosis, necrosis, lesion) were read while several molecular labels were not. The composition of the graph thus also records what the reading stage saw. For a user this is an argument for running one slide at a time and merging the graphs, which the stage supports, rather than for drawing everything on one slide.

#### 4.6.0.4 The DUX4 use case and the nature of the bridges

The DUX4–HMGB1 interaction assertion is the kind of connection a reader of the deck is looking for: the danger signal the researchers placed downstream of mitochondrial damage is, in RTX-KG2c, a physical interaction partner of DUX4, found in a co-purification screen of DUX4-associated proteins in muscle cells (Ansseau et al., 2016), a connection that the slide does not draw; it ties the nuclear defect of FSHD to the danger signal that the mitochondrial pathway places downstream of oxidative damage, with a primary source (IntAct) the user can open. The other paths of Figure 4d are weaker, and the figure shows why: in RTX-KG2c under this profile, the cheapest route between a transcription factor and a signalling pathway often runs through a shared cellular location (*expressed in neural nucleus*, *located in cytosol*), an edge that is true, sourced and nearly uninformative. The effective weights of these edges under the biochemical profile (0.33 for *located in*, 0.67 for *expressed in* and *colocalizes with*, 1.0 for hierarchical edges) are higher than that of the one assertion a biologist would retain, the IntAct interaction (0.05, the product of the predicate weight and the 0.3 weight given to the source), which wins only because it is a single hop. This argues for exposing the predicate and source of every edge in the interface, as the local console does, rather than for suppressing localisation edges, since co-localisation is also the first thing a bench scientist checks before testing an interaction; it also argues for a source weight for curated interaction databases closer to that of pathway databases.

### 4.7 Limitations and future directions

#### 4.7.0.1 Scope of the evaluation

The evaluation has the limits of a single-team study. The expert gold of the decks and the audit of the benchmark were each produced by one expert, without a measure of inter-annotator agreement. The decks come from one research team and one disease area, and five of the six decks with an expert gold are in French while the description is written in English, which lowers their exact recall and their typing scores. The pathway signatures of the case study were written after the graphs were built; they are a reading aid, not a validation of the hypotheses drawn on the slides. Finally, we are not aware of another end-user tool that starts from slides and returns typed, linked and contextualised entities, and we therefore compared each stage with its own baselines (retriever alone, hosted models, fully local linking) rather than the whole workflow with another system.

#### 4.7.0.2 Running time

Most of the wall-clock time goes to the typing agent (69 to 78% of a run, 15 to 25 s per candidate mention), so a one-slide deck takes half an hour and a two-slide deck one hour on the test laptop. End users should therefore treat BiomedCAT as a job to submit and return to rather than as an interactive query tool; the local console queues documents and runs them one after the other for that reason. Two levers for reducing this time are discussed below.

#### 4.7.0.3 Encoder models and distillation

First, a natural question is whether the typing stage needs a generative model at all, or whether a biomedical encoder (PubMedBERT, SapBERT, BioLinkBERT) run locally would do the same work in milliseconds. Our results answer the zero-shot half of the question. An encoder used by similarity, embedding the mention and the Biolink class descriptions and exemplars and taking the nearest class, corresponds to the retriever-only configuration of BiomedCAT: it reaches 0.42 on the audited benchmark against 0.71 for the agent built on top of it (Figure 2B.b), and substituting a biomedical encoder for the default one does not close the gap (hit@1 on the public slice: pubmedbert-base-embeddings 0.35, the default clinical ModernBERT 0.33, SapBERT 0.29, Supplementary Table S6, E11b). The 28 points gained by the agent come from choices that similarity cannot make: between a class and its parent given the sentence, between an entity and the process it is involved in, and the abstention (NONE) for strings that name nothing or name a person, which the privacy rule requires and which the agent returned on all four personal-data terms of the audit.

Second, a fine-tuned encoder classifier over 147 classes needs a labelled corpus at Biolink granularity, and none exists: the synthetic labels are wrong or ambiguous for 55% of the audited mentions, and the expert gold amounts to 664 synthetic mentions and some 300 deck terms, with many classes under ten examples. Two routes follow from this work. The first is distillation: typing a large synthetic corpus with the agent, keeping the verdicts that satisfy the audit rules, and training a small encoder on them, keeping the agent only for abstention and for the mentions the encoder scores as ambiguous; the per-term outputs deposited with this article are the seed of such a set. The second is a zero-shot span extractor of the GLiNER family for stage 1, where the recall-first prompt is the simplest part of the pipeline to replace with a model a hundred times faster. Both would move most of the typing time, currently 15 to 25 s per mention, to a model of a few hundred megabytes, while leaving the contextual decisions to the agent.

## Author Contributions

SM: Conceptualization, Methodology, Software, Validation, Formal analysis, Investigation, Data curation, Writing – original draft, Writing – review & editing, Visualization, Supervision, Project administration. SC: Methodology, Software, Data curation, Writing – original draft. LH: Writing – review & editing. LT: Writing – review & editing. JD: Methodology, Investigation, Data curation, Writing – review & editing. CH: Conceptualization, Methodology, Investigation, Data curation, Writing – review & editing.

## Acknowledgments

We thank the CATRAX team of the NIH/NCATS Biomedical Data Translator consortium for providing the knowledge-graph resources (RTX-KG2) and the ARAX and Plover services used in this work, and Gaurav Vaidya (RENCI) for his work on Babel, the Name Resolver and the Node Normalizer, on which the linking stage relies. We thank the Ensimag school of Grenoble INP^†^ and the teaching team of the MOSIG master’s programme for making the student collaboration possible, and in particular Prof. Grégory Mounié (Univ. Grenoble Alpes, CNRS, Inria, Grenoble INP^†^, LIG, 38000 Grenoble, France) for his feedback at the students’ project presentations. We are grateful to the members of the DAISY team of LIG for discussions during the development of the method.

## Supplementary Material

The Supplementary Material for this article (Supplementary Methods, Tables S1 to S10 and Figure S1) is available as a separate file accompanying this preprint.

## Data Availability Statement

The source code of the BiomedCAT pipeline (reading, typing, linking, context graph and local console) is available at https://github.com/AI4Health-DAISY-LIG/BiomedCAT (release v1.0.0). The synthetic typing benchmark is deposited on Hugging Face at https://huggingface.co/datasets/sandrine-muller/biolink-typing-benchmark (https://doi.org/10.57967/hf/10796, version 1.0): its public slice of 1,075 mentions with the original labels, the expert audit (verdict and corrected class) and the canary string. Its generation, verification, split and evaluation scripts and the baseline typers are at https://github.com/AI4Health-DAISY-LIG/KG_nodes_eval. The private slice is withheld by design (Methods) and can be obtained from the corresponding author for the evaluation of future models, under a no-redistribution agreement. The source data of every figure and table are deposited on Zenodo as two versions of the same record, the source-data archive (https://doi.org/10.5281/zenodo.23197942) and an addendum (https://doi.org/10.5281/zenodo.23201488). The source-data archive holds the run of each slide deck (reading, typed mentions, identifiers, context graph, per-second memory and GPU samples), the harmonized expert gold of the decks with its verification against the curation sheets, the scores of every run against it, the public slice of the typing benchmark with its expert audit and the audited scores of every typing system, the recorded hardware, software and remote-service versions, and the data file behind each figure (the archive mirrors the data/publication/ layout of the repository; MANIFEST.json lists the SHA-256 of every file). The same archive holds the retrieval exemplars (3,712 mentions) and the input documents of the article (the seven slide decks and the image, with their metadata sheet). The addendum holds the context graphs of the FSHD deck under each of the six profiles, with the corresponding presence checks, and the context graphs behind Figures 3 and 4 (the three pathway slides, their integration and the summary slide of the FSHD1 inflammation deck).

The pipeline runs locally; a Docker configuration is provided. The scispaCy model en_core_-sci_sm is optional (SCISPACY=0 runs without it, which is the configuration of every published number).

## Generative AI statement

The method, the study design and the original codebase were conceived and developed by the authors. Generative AI tools were used at four levels, all under the authors’ direction and review: (i) as components of the method itself (local language models for OCR, entity typing and linking, as described in Methods); (ii) to generate the synthetic typing benchmark (gpt-oss-120b) and the retrieval exemplars (mistral-small), whose labels were then audited by the authors (Section 2.5.2.3); (iii) as a coding and analysis assistant (Claude and Gemma) for the implementation of retrieval and evaluation components, test suites, experiment scripts, result tables and figures, the user interface and deployment configuration; and (iv) as a writing assistant (Claude) for drafting and editing parts of the manuscript and of the Supplementary Material, which the authors revised. All AI-assisted code, analyses and text were reviewed by the authors, and every reported number was verified against the raw outputs; the expert ground truth and the interpretation of the results are the authors’ own.

