## Supplement information for "BiomedCAT: Biomedical Context Analysis Tool for Supporting Mechanistic Hypothesis Generation via User-defined Contextual Subgraph Extraction"

*Supplementary Material of a preprint that has not been peer reviewed, made available under a CC BY 4.0 licence.*

### 1 Supplementary Methods

#### 1.1 Profiles shipped with BiomedCAT

Each profile is a small set of anchor classes with a priority (1, primary; 0.5, secondary context); the scope, the predicate weights and the category weights are derived from them as described in the main text (Methods, *Automated construction of scopes from end user profiles*).

- *biochemical actions*: molecular-level mechanisms: binding, interaction, regulation, and pathway participation. Anchored on *gene*, *gene family*, *transcript*, *polypeptide*, *protein family/domain*, *regulatory region*, *coding sequence*, *sequence variant*, *genotype*, *haplotype*, *genome*, *macromolecular complex*, *nucleosome modification*, *chemical entity*, *biological process or activity*, *disease or phenotypic feature*, *organismal entity* and *exposure event*.

- *pharmacological intervention*: drug-discovery and pharmacology angle: a chemical entity, the gene or protein it targets, its mechanism of action, and its clinical use. Anchored on *chemical entity*, *gene*, *polypeptide* and *macromolecular complex*, *disease or phenotypic feature*, *clinical entity* and *biological process or activity* (priority 1), with *organismal entity*, *exposure event*, *transcript* and *reagent targeted gene* as secondary context (priority 0.5).

- *clinical mechanisms*: organism and patient level: disease, phenotype, predisposition, treatment and trial evidence. Anchored on *disease or phenotypic feature*, *clinical entity*, *gene*, *polypeptide*, *chemical entity*, *organismal entity*, *exposure event*, *activity*, *attribute* and *physical entity*.

- *epidemiological risk*: population and public-health angle: exposures, risk factors and disease occurrence across individuals and populations, with environmental and demographic context. It does not include the molecular detail of the mechanism nor the single-patient course of care. Anchored on *exposure event*, *organismal entity*, *disease or phenotypic feature*, *planetary entity*, *attribute*, *activity* and *study result* (priority 1), with *gene*, *chemical entity*, *biological process or activity*, *evidence type* and *clinical entity* as secondary context.

- *genetic determinants*: heredity and molecular-genetics angle: the causal variant, its mode of inheritance, and the gene or regulatory region it alters. It does not emphasize what the alteration does downstream nor how it presents in the clinic. Anchored on *gene*, *sequence variant*, *genotype*, *haplotype*, *genetic inheritance*, *genome*, *regulatory region*, *coding sequence*, *exon*, *nucleic acid sequence motif*, *nucleosome modification*, *reagent targeted gene* and *disease or phenotypic feature* (priority 1), with *organismal entity*, *transcript*, *polypeptide* and *biological process or activity* as secondary context.
- together with a *uniform* baseline (every entity branch and predicate at weight 1, undirected traversal, no source or knowledge-level weighting).

#### 1.2 Exemplar index of the retriever

The typing agent's retrieval index combines two complementary sources of evidence for each Biolink class: the class's formal definition and hierarchical position in the Biolink model, and a set of short surface-form exemplars, realistic mentions that a document might actually contain. Definitions capture what a class means; exemplars capture what it looks like in text, which a definition written in ontological language often does not ("DM" or "type 2 diabetes mellitus" for the class *disease*). A query is matched against both collections and the two rankings are fused by reciprocal rank fusion.

**1.2.0.1 Published index (v4)** Exemplars were generated class by class with `mistral-small-3.2-24b-instruct` (30 requested per class) in four surface forms, canonical name, multi-word compound, abbreviation or symbol and orthographic variant, under the same leakage rules as the benchmark: an exemplar that repeated the class name, that duplicated an exemplar of another class or that coincided with a mention of the typing benchmark was rejected (722 rejections: 328 benchmark overlaps, 324 duplicates or class names, 70 length or form). The index holds 3,712 exemplars for 145 classes (1,174 canonical names, 1,023 compounds, 774 abbreviations, 741 variants), each indexed as its own document carrying its class (`data/biolink_exemplars.parquet`, deposited with the article). Adding this leg improved the retriever hit@1 by 7 points (Table S6, E1).

**1.2.0.2 Variant drawn from the knowledge graph (v5, not retained)** An alternative index drew the exemplars from the names and recorded synonyms of RTX-KG2c nodes, so that every exemplar corresponds to an entity present in the graph. For each Biolink class with a matching KG2c category, up to 30 nodes were sampled, with a bias towards better-connected nodes (number of edges) so that broadly recognised concepts ("type 2 diabetes mellitus", "Crohn disease") are preferred over narrowly documented ones; each node contributed its canonical name and one or two synonyms. Of the 157 non-root Biolink classes retained for typing, 52 map to a category instantiated in the KG2c build; the other classes kept their generated exemplars. The variant totals 3,968 mentions over 146 classes. It lowered the retriever hit@1 by 2 points (Table S6, E2): KG2c names stay close to the identifiers and formal labels that scientists rarely use in their prose, and the variant was abandoned.

##### 1.3 Synthetic typing benchmark: construction, split and audit

This section gives the protocol behind the benchmark summarised in the main text (Methods, "NER task") and the figures needed to reproduce every subset used in the article. Files and scripts are in the companion repository `KG_nodes_eval` (Data Availability Statement).

###### 1.3.1 Generation

Classes were read from the Biolink model release 4.4.4. The two structural roots (`entity`, `named thing`) were excluded; every other class, mixins included, is a stratum with a quota of ten mentions. For each class the generator (`gpt-oss-120b`, seed 77) received the class name, definition, examples, parent, siblings and children, and was asked for real biomedical entities of one to four words written as they appear in papers or slides, with an explicit quota of four surface forms: canonical name, abbreviation or symbol, orthographic variant (plural, casing, hyphenation, common misspelling) and multi-word compound. A mention was rejected when it repeated the class name, one of its examples or aliases, or any word of five letters or more of its definition; when it duplicated a mention already accepted for another class (normalised string); or when the verifier (`deepseek-v4-flash`), asked to choose among the target class, its parent, its siblings and its children, did not return the target. The verifier's choice is stored with every mention (`verifier_choice`). 895 mentions were rejected; the rejection log is released with the benchmark.

###### 1.3.2 Composition

1,356 mentions, 147 classes. Ten of the 157 non-root classes kept no mention, the verifier assigning all of them to a child or sibling class or to none: *anatomical entity*, *biological process or activity*, *chemical entity*, *disease or phenotypic feature*, *exposure event*, *molecular mixture*, *evidence*, *retrieval source*, *journal article* and *book chapter*. The classes that reached fewer than ten mentions are listed in the released statistics file. Surface forms: 412 canonical, 382 abbreviation, 282 compound, 280 variant. Depth in the class tree: 144 (depth 2), 620 (3), 340 (4), 215 (5), 37 (6).

###### 1.3.3 Public/private split and canary

The benchmark was split with seed 2026, stratified by class and surface form, with 20% of each class held out (at least one mention when the class has three or more). Public slice: 1,075 mentions, 147 classes (340 canonical, 314 abbreviation, 211 variant, 210 compound; depths 2 to 6: 114, 492, 271, 169, 29). Private slice: 281 mentions, 146 classes (72 compound, 72 canonical, 69 variant, 68 abbreviation; depths 2 to 6: 30, 128, 69, 46, 8); one class (`phenotypic sex`) has public mentions only. A canary string, `BIOLINK-TYPING-BENCHMARK CANARY GUID` followed by a unique identifier, is written in the schema metadata of the public Parquet file, as a field of every line of the public JSONL file and in a `CANARY.txt` file. If a model asked to complete the prefix reproduces eight or more hexadecimal characters of the identifier, it has seen the release in training and must be scored on the private slice. Scores in this article are reported on the public slice; the private slice is reserved for future models and is not deposited anywhere.

##### 1.3.4 Reference subset

A fixed subset of 300 public mentions (seed 77) served for every configuration decision of Table S6. Its binomial sampling error is about 2.8 points; the run-to-run variation of the agent on the full benchmark is about 1 point.

##### 1.3.5 Expert audit

The labels of 664 distinct public mentions were judged against the Biolink class definitions by the bioinformatician expert, in two passes (18 and 22 September 2026). Mentions were selected in five groups, in this order: (1) the 300 reference mentions; (2) 133 mentions outside the reference subset on which three independent typers (Claude, DeepSeek, gemma-3-1b) agreed on another class than the label; (3) 83 mentions completing the class coverage; (4) in the second pass, 43 reference mentions kept "ok" in the first pass but contradicted by a run not displayed in the first pass, re-judged with the same care (the second verdict supersedes the first); (5) 148 public mentions added so that every reading scope (coarse stratum) reaches the same number of scorable mentions, taken in increasing order of ambiguity risk within the form mix of the stratum. Model outputs were shown as a triage hint only; the instruction was to judge the label, not the models. Verdicts: *ok*, *wrong* (with the class the expert would assign), *ambiguous* (several classes defensible for the mention as written). Domain and range constraints were not used, because the first pass found them missing or too narrow for several classes. Results: reference subset, 113 ok, 125 wrong, 62 ambiguous; all 664 audited mentions, 298 ok, 216 wrong, 150 ambiguous (audit file of 4 October 2026; rescoring rerun on 6 October). The released file carries, for every public mention, the original label, whether it was audited, the verdict, the corrected class when the label was wrong, and a derived `gold_class` column (the original label when confirmed, the expert class when wrong, empty otherwise).

#### 1.4 Reading-stage factor test on the FSHD deck

The reading stage does not transcribe a slide: a vision–language model describes it as a numbered list of dense statements, and the entities are extracted from these statements downstream. We measured how the description of the two slides of the FSHD deck (Zenodo, `https://doi.org/10.5281/zenodo.23197942`) depends on three factors, the pipeline's own prompt being kept: the model (`gemma4:e4b-it-qat`, the pipeline's classification model, or `qwen2.5vl:3b`, its reading model), the temperature ($T = 0$ and $T = 1$ for Gemma, $T = 0$ for Qwen) and the profile that parameterises the prompt (none, i.e. the bare prompt; *biochemical actions*; *genetic determinants*; the last two add to the prompt their list of entity kinds to name and their reading focus, with no custom text). Since the pipeline sends a fixed random seed, the $T = 1$ conditions were run with two seeds. This gives 12 conditions and 24 descriptions (Supplementary Table S4). Each description was read against its slide by the first author; an automated check of the elements printed or drawn on each slide (24 on slide 1, 21 on slide 2; Supplementary Table S5) is reported as a descriptive count only, since it tests the presence of a token, not the correctness of the statement that contains it.

##### 1.4.1 Design

Script `tests/ocr_factors_fshd_test.py` (no change to the pipeline code) rasterises `Dataset/FSHD.pdf` with the pipeline's rasteriser, builds the reading prompt with `biomedcat.prompts.ocr_description` from the entity scope and reading focus of the selected profile, and calls Ollama through `biomedcat.runtime.chat` with the pipeline's options (8,192 new tokens, hidden reasoning off, `keep_alive` 0). Factors: model $\in$ {`gemma4:e4b-it-qat`, `qwen2.5vl:3b`}; temperature $T \in \{0, 1\}$ for Gemma, $T = 0$ for Qwen, seeds 0 and 1 at $T = 1$; profile $\in$ {none, *biochemical actions*, *genetic determinants*} as shipped in `data/profiles`. The bare prompt is 2,145 characters; the biochemical profile adds 30 entity kinds and a 281-character focus (2,910 characters in all), the genetic profile 15 kinds and a 310-character focus (2,683). Run on 4 October 2026.

##### 1.4.2 Model

At $T = 0$ the two models fail differently. Qwen reproduces the printed text with high confidence: the three sentences of slide 1, the gene symbols (SMCHD1, DNMT3B) and, alone among the twelve conditions, the tokens *D4Z4*, *FSHD1* and *FSHD2* exactly as written. Gemma corrupts short tokens in every one of its $T = 0$ descriptions (*D4Z* for D4Z4, *FSD1*/*FSD2* for FSHD1/2, *F-S-H-D*, *DAD* for TAD), and garbles the reference line it is the only model to read ("Beneri CRS and Zamm PB"). Conversely, Gemma reads the annotated schematic of slide 2 more completely: without a profile it names the DNA-binding domains HD1 and HD2, the N- and C-termini, the TAD, the DUX4 target gene and the five outcomes (15 of 21 elements), and it gives the only defensible reading of the FSHD section ("replacement by fibrotic/inflammatory tissue"). Qwen, without a profile, names none of the domain labels (10 of 21), rewrites the five parallel arrows as a serial chain (DNA binding → aggregation → impaired NMD → impaired myogenesis), and, when the prompt asks for colours, invents them: a blue/red/grey code for FSHD1, FSHD2 and control on slide 1, and "red indicating normal tissue and black highlighting areas of damage" on an H&E section. Qwen also mirrors the prompt's directives as section-like statements ("The structural elements include...", "The biochemical markers are...", "chromosomes and RNA loops are not explicitly shown") and closes with the summarising sentences the prompt forbids. On the same machine, reading a slide took 17–22 s with Gemma and about six times longer with Qwen.

Two errors are shared by all conditions. On slide 1 every description assigns FSHD1 to the unstable DUX4 mRNA and FSHD2 to the stable one, a pairing the slide does not make (both mechanisms point to the array; the stable transcript is the one that ends with the "(A)n" tail), and none reads "(A)n" as polyadenylation (Gemma: "amplification", "labeled 'a' and 'n' "). On slide 2 no description mentions the fluorescence panels, and only one (Gemma, genetic profile) infers the mitochondrion behind the ROS symbol.

##### 1.4.3 Temperature

Raising Gemma to $T = 1$ did not add content and degraded name fidelity. Over the twelve $T = 1$ descriptions the invented or corrupted names include *DNMT3A* for DNMT3B, *DUSH gene*,

*Duplexin*, *Tandem Array Domain* and *NME (Non-Mendelian Elimination)* as expansions of TAD and NMD, *F5H1*, *Facioepiglottic*, and, in two of the six readings of slide 2, the wrong disease altogether ("Duchenne muscular dystrophy", "dystrophin"). Formatting drifted (LaTeX arrows and `\text{}` macros, bold Markdown), which matters because the extractor downstream keeps mentions as written. The two seeds of the same condition disagree on substance (one names Duchenne, the other does not), so the variance is real and not a seed artefact. The `[UNCLEAR]` marker, requested by the prompt, was used 14 times at $T = 1$ and never at $T = 0$ by either model. Element counts at $T = 1$ were in the same range as at $T = 0$ (slide 1: 16–21 vs 17–18; slide 2: 12–14 vs 11–15).

##### 1.4.4 Profile

The profile changed the register more than the coverage. With the *genetic determinants* profile Gemma writes of the "hereditary basis", a "regulatory region", a "genetic/epigenetic alteration" and, on slide 2, of "a specific genetic variant or regulatory element within the DUX4 locus" that the slide does not show; with *biochemical actions* it writes of "the anatomical entity of skeletal muscle fibers" and of "a polypeptide structure". These phrases take their words from the entity list and the reading focus injected into the prompt. Qwen with a profile added the domain labels it had missed without one (HD1/HD2 with both profiles; N-/C-ter and TAD with *genetic*), but also the prompt-echoing statements noted above and, with *biochemical*, "RNA loops that interfere with NMD", an example from the prompt turned into a finding. On the element counts the profile had no consistent direction: Gemma at $T = 0$ scored 18/17/18 on slide 1 and 15/11/14 on slide 2 for none/biochemical/genetic; Qwen 20/21/21 and 10/13/11.

##### 1.4.5 Errors shared by all conditions

- Slide 1: FSHD1 paired with the unstable transcript and FSHD2 with the stable one (12/12); "(A)n" never read as a poly(A) tail.
- Slide 2: fluorescence panels never described (0/12); the mitochondrion inferred once (Gemma, $T = 0$, genetic); the sequence logo read once, partially (Qwen, no profile: "TAATCA").

##### 1.4.6 Corrupted or invented names (Gemma)

- $T = 0$: D4Z (for D4Z4), FSD1/FSD2 (for FSHD1/2), F-S-H-D, DAD (for TAD), "Beneri CRS and Zamm PB" (for Banerji & Zammit)
- $T = 1$, in addition: DNMT3A, F5H1/F5H2, FHSD, FHS, Facio-cophumeral, FacioCapuholumuscular, Facioepiglottic, Fibroscapulohumeral, DuX4, DUSH gene, Duplexin, Tandem Array Domain (TAD), NME (Non-Mendelian Elimination), chromosome 4G, hemimethylation, "Duchenne muscular dystrophy" and "dystrophin" (two descriptions of slide 2), "immunohistochemical staining for ROS and apoptotic markers".

##### 1.4.7 Prompt echoes and invented visual attributes (Qwen)

- "blue for FSHD1, red for FSHD2, and gray for control" (slide 1, biochemical);
- "red indicating normal tissue and black highlighting areas of damage" (slide 2, none);
- "RNA loops that can interfere with NMD" (slide 2, biochemical);
- "The structural elements include... / The biochemical markers are... / The morphological features include... " (slide 1, genetic);
- closing summary paragraphs in four of six descriptions.

#### 1.5 Selection of the typing configuration

The configuration of the Biolink typing step (retriever, agent and model parameters) was selected through twenty-one experiments, summarized in the main text and listed in Table S6. Each experiment changed one element relative to the configuration current at that point and was stated beforehand as an observation, a hypothesis and an acceptance rule. The experiments are listed in the order in which the reasoning unfolded; rows E10, E14 and E15 refer back to earlier rows.

All agent experiments used `gemma4:e4b-it-qat` served by Ollama with greedy decoding (temperature 0, seed 0), unless the experiment itself changed this parameter. Two test sets were used. Retriever experiments were scored by hit@1 on the 1,075 public terms of the synthetic benchmark (the public split of the 1,356-term benchmark). Agent experiments were scored by the accuracy of the final Biolink class on a fixed subset of 300 reference terms drawn from the same benchmark and compared with the original labels produced by the generator ("original gold").

A change was considered an improvement only if it exceeded 5 points. This margin lies above the binomial sampling error of a 300-term subset (about 2.8 points) and above the variation between three complete runs of the agent on the 1,356 terms ($0.445 \pm 0.010$). The expert audit of the benchmark labels (main text, Methods, "Synthetic curated set") raised the absolute accuracy of every system. The figures of Table S6 are therefore to be read as differences between configurations rather than as absolute performance: on the 300-term subset, the agent scores 0.410 with the tool loop and 0.470 with the single-call prompt introduced in E14.

Experiments E1 and E2 concern the exemplar index, which is described in Section 1.2.

#### 1.6 Evaluation on the slide decks

##### 1.6.1 Resource measurement

Time per stage is read from the run log of each deck (`OCR done`, `NER done`, `Norm done`, `Context graph done`) and the total from the run record. Memory was measured by a wrapper (`scripts/measure_pipeline_resources.py`) that starts the pipeline in the published configuration and samples once per second the resident set of the pipeline process tree, the resident set of the Ollama server processes, the system memory in use, the GPU memory in use (`nvidia-smi`) and the model resident in Ollama. Samples are attributed to stages by the timestamps of the log markers; every figure is the peak within the stage, relative to the idle level sampled before launch. The reported RAM is the system memory in use above idle, which includes the driver-backed buffers that mirror GPU allocations on Windows and is therefore the quantity a machine without GPU would need in addition to the weights. Per-second samples are deposited with each run (`resources_samples.csv`).

**1.6.1.1 Energy estimate** Energy was not metered. It is bounded by the wall-clock time of each run multiplied by the power draw of the laptop, between 60 W (CPU load, GPU idle) and 200 W (GPU at full load): 0.03 to 0.14 kWh for a one-slide deck (33 to 42 minutes) and 0.06 to 0.20 kWh for the two-slide FSHD deck (59 minutes). The carbon footprint uses the greenhouse-gas intensity of electricity generation in 2025 reported by the French transmission system operator RTE (*Bilan électrique 2025*): 19.6 g $CO_2$e/kWh for France and 178 g $CO_2$e/kWh for the European average (European Union, Switzerland, Norway and United Kingdom), that is 0.6 to 3.9 g and 6 to 35 g of $CO_2$e per run.

##### 1.6.2 Expert gold of the slide decks

**1.6.2.1 Harmonization** The curated spreadsheets of the seven decks (one per deck, different layouts) were merged into one file by `scripts/build_gold_harmonized.py` without changing any curated value: the 3,891 copied cells were re-read from the sources and compared, with no difference. Each row carries its deck, source file and sheet, a kind (*positive*: a term written on the slide with a Biolink class; *implicit*: expected by the expert but not written; *adversarial*: must receive neither class nor identifier; *na_on_slide*: written on the slide but judged not to be an entity; *removed*), the explicit or implicit flag (from the sheet when it has one, otherwise assumed explicit because the drafts list terms read on the slide with their slide zone), the curation status and the curator's note. Two derived columns are added: a compound flag and the KG2c label of the gold identifier, used only to match English readings of French terms. Identifiers are not scored. Counts: FSHD 32 explicit and 7 implicit positives, VoieA 42, VoieB 42, VoieC 34, VoiesA-C 60, FSHD1multiscales 68 explicit and 1 implicit, glycolysis 8 explicit and 11 implicit; 143 adversarial terms in all.

**1.6.2.2 Scoring rules** A gold term is matched through up to three forms: the term as written, its English query when the sheet has one, and the KG2c label of its identifier. *Reading, exact*: one form, lower-cased and accent-stripped, is a substring of the text produced by the reading stage. *Reading, lenient*: every token of one form (three characters or more, stop words removed) has a token in the reading that is equal or shares its first five characters, a deliberately crude stemmer that accepts French/English cognates (*mitochondrie*/*mitochondria*, *apoptose*/*apoptosis*) and plurals. *Extraction*: an extracted mention equals one form, or contains it, or is contained in it, with the same token tolerance. *Typing*: on found positive terms with a gold class, the class of the matched mention equals the gold class, or lies at hierarchical distance one in the Biolink tree (parent or child). Reading recall is reported separately for explicit and implicit terms; typing is reported on explicit terms only, because implicit terms have no surface form to match. Per-term verdicts are deposited (`gold_eval/per_term.csv`).

##### 1.6.3 Linking: online services against a fully local mode

On the FSHD deck the two deployment modes share the reading and typing (extraction recall 0.91, typing accuracy 0.48 on the found terms) and differ only at the linking stage (Supplementary Figure S1). Of the 29 expert terms found, 6 have no identifier in RTX-KG2c or Node Normalizer (D4Z4 array, repeats, impaired myogenesis, DNA binding domains, DUX4 binding site, DUX4 target gene) and are excluded. On the 23 others, the online mode assigns the expert identifier to 65% under each of three nested criteria (same CURIE, same canonical RTX-KG2c node, same Node Normalizer clique); the fully local mode reaches 26%, 30% and 26%. Online, 70% of the 175 extracted mentions receive an identifier, right or wrong, against 42% offline. The loss is concentrated on acronyms and on common anatomical names absent from the KG2c name table. Linking is the only stage where leaving the machine buys accuracy; the user chooses it per run.

##### 1.6.4 Multi-scale case: pathway signatures, structure and paths

The five slides (`VoieA.pdf`, `VoieB.pdf`, `VoieC.pdf`, `VoiesA-C.pdf`, `FSHD1multiscales.pdf`; FSHD1 inflammation deck v13) were run on 4 October 2026 with the published configuration (reading `qwen2.5vl:3b`, typing and linking `gemma4:e4b-it-qat`, public resolvers; `data/eval/master_results`, sheet *pipeline_runs*). Each run's linked entities seeded a context graph with the biochemical profile, 2 hops, hub cap 500, per-category cap 300, score floor at the median of each hop, $\alpha = 0.5$, prediction edges kept. The analysis script (`scripts/voie_multiscale_case.py`) reads the exported `nodes.csv`, `edges.csv` and `summary.json` of each graph and writes `data/eval/voie_multiscale/`; the figures come from `scripts/make_voie_panel.py`.

**1.6.4.1 Reading of the slides** Mentions, linked mentions and distinct seeds: slide A 103 / 77 / 20 (9 Biolink types among linked mentions); slide B 88 / 63 / 19 (11 types); slide C 79 / 55 / 24 (12 types); integrated slide 64 / 39 / 20 (9 types); summary slide 93 / 69 / 25 (17 types). Linking errors propagate to the seeds: the integrated slide carries *pulmonary immaturity* (MONDO:0002929, linked from "regeneration immaturity"), *NS1 mediated effects on host pathways* (Reactome, from the bare word "pathways"), *Expression of SOCS3* (from "myonuclear expression") and *inflammatory myopathy with abundant macrophages* (from "macrophages/FAPs association") among its seeds; slide B carries *neurodegeneration, childhood-onset, with multisystem involvement due to mitochondrial dysfunction* (from "mitochondrial dysfunction") and slide A *11q aberration* (from "transcriptional aberration"). They were kept as the pipeline produced them; none of the signature nodes depends on them, and they illustrate the linking failure mode of the judge on generic or compound mentions discussed in the Results.

**1.6.4.2 Pathway signatures** Table S7 lists the ten patterns of each signature; Tables S8, S9 and S10 give the structure of the five graphs, the recovery of the signatures and the distances between them. A pattern is matched case-insensitively as a substring of the RTX-KG2c node name, or as the whole name when prefixed by "=" in the script. A signature node is counted present when at least one kept node matches; its rank is the rank of the best-scored match inside its Biolink category (seeds are not ranked). No signature node is a seed of any of the five decks.

###### 1.6.4.3 Best-weighted paths from DUX4

The path cost reproduces the stage's `path_-weight`: the sum over edges of $-\log w$ plus, for every node, $\log\log_2(2 + \text{degree})$ with the degree taken in RTX-KG2c (column `degree` of `nodes.csv`), the two endpoints contributing a constant. Without the degree term, weight-1 edges (`subclass_of`, `manifestation_of`) are free and the best path wanders through ontology hierarchies for tens of hops. The paths are read below with predicate, primary source and profile weight. In the slide-A graph, DUX4 *expressed in* neural nucleus (NCIT, 0.67), which DHX15 is also *expressed in* (NCIT, 0.67), DHX15 being a participant of the antiviral innate immune response (*has participant*, GOA, 0.20). In the slide-B graph, DUX4 *physically interacts with* HMGB1 (IntAct, knowledge assertion, 0.05); the same edge is the DUX4-to-B path in the slide-C graph. In the slide-C graph, DUX4 is a participant of the Cell Differentiation process (*has participant*, NCIT, 0.20), *subclass of* cellular developmental process, *subclass of* cellular process (GO, 1.0), of which the execution phase of apoptosis is a subclass (GO, 1.0). In the A+B+C graph, the path to signature A runs as in slide A with RELA in place of DHX15; the path to signature B goes from DUX4 *located in* cytosol (GOA, 0.33), where NLRP3 is also *located* (GOA, 0.33); the path to signature C is that of slide C. In the summary-slide graph, DUX4 reaches the DDX58/IFIH1-mediated induction of interferon-$\alpha/\beta$ through neural nucleus, APP and its transcript APP-210 (*transcribed from*), and the collagen-containing extracellular matrix through cytosol and TGM2 (*colocalizes with*, GOA, 0.67). The node and edge tables of the three windows of Figure 4 (the full windows, of which the figure draws the largest connected component) and the edges of every path are regenerated by `scripts/make_fig4_dux4.py` of the BiomedCAT repository; the figure itself was laid out in Cytoscape from these tables.

###### 1.6.4.4 DUX4 target-activation reactions

Ranks inside the kept MolecularActivity category (300 nodes): slide A 13–27, slide B 8–16, slide C 15–26, integrated slide 12–20, summary slide 18–27; all six reactions present in every graph (`scripts/fshd_dux4_mechanism_case.py` gives the same table for the six profiles of the FSHD deck).

###### 1.6.4.5 Limits

The signature lists were written by us from the slides and from the Reactome and GO vocabularies after the graphs were built; a signature node absent from a graph may be absent from RTX-KG2c under the biochemical profile's scope rather than unreachable. All five graphs use the same profile; the comparison of profiles is not repeated here. The paths of Figure 4d of the main text are the best-weighted paths of the kept graph, which is already pruned; a path in the full RTX-KG2c may be shorter.

##### 1.6.5 Compounds tested in FSHD

The list holds 21 compounds that have been tested in people with FSHD, compiled from Clinical-Trials.gov and published trials (`data/eval/fshd_drugs_presence.json`): ten investigational agents (DUX4-targeting oligonucleotides, epigenetic editing, anti-myostatin antibodies, p38 inhibition and others), eight marketed drugs tested for repurposing and three supplements or supplement combinations. Each compound was mapped by name to its RTX-KG2c node; combinations were matched on one component (folic acid; ascorbic acid), somatropin on the GH1 gene, which is the only node of growth hormone in the graph, and testosterone enanthate on its RTX-KG2c clique (named *testosterone cypionate*). The context graphs were rebuilt from the publication run of the FSHD deck (5 October 2026) under each of the six profiles, with the parameters of the case study, by `scripts/build_fshd_profile_graphs.sh`; the stage's `-find` option records, for a compound reached but not kept, its hop and its rank among the reached nodes of its category. `scripts/graph_presence.py` writes the presence table (`data/eval/fshd_presence_pub.json`) and `scripts/fshd_drugs_venn.py` its Venn diagram. The rebuilt biochemical graph keeps the same three compounds as the deposited graph of the publication run; the two differ by 196 nodes, all tied at the category cap (Methods, *Context graph construction*).

### 2 Supplementary Tables

#### 2.1 Per-deck results

Table S1: Reading recall (exact / lenient) and typing accuracy (exact class / within one step) on the explicit expert terms of each deck; $n$ = explicit terms, $n_t$ = found explicit terms with a gold class. Values of the publication runs of 5 October 2026 (`data/publication/gold_eval/summary.csv`).

| Deck | $n$ | reading exact | reading lenient | extraction | $n_t$ | typing exact / ≤1 step |
|---|---|---|---|---|---|---|
| FSHD (EN, 2 slides) | 32 | 0.81 | 0.94 | 0.91 | 29 | 0.48 / 0.66 |
| VoieA (FR) | 42 | 0.62 | 0.71 | 0.62 | 26 | 0.19 / 0.31 |
| VoieB (FR) | 42 | 0.31 | 0.52 | 0.62 | 26 | 0.35 / 0.54 |
| VoieC (FR) | 34 | 0.59 | 0.79 | 0.77 | 26 | 0.19 / 0.42 |
| VoiesA-C (FR) | 60 | 0.28 | 0.48 | 0.47 | 28 | 0.18 / 0.54 |
| FSHD1multiscales (FR) | 68 | 0.35 | 0.59 | 0.54 | 37 | 0.30 / 0.49 |
| Glycolysis (image) | 8 | 0.25 | 0.25 | 0.50 | 4 | 0.00 / 0.50 |

#### 2.2 Existence gate on the gold suite

Table S2: Existence gate on the FSHD gold suite v2: 32 expert terms and 34 invented terms (nonsense words, absurd compounds, near-misses of real symbols, author names and years), submitted to the gate alone, without typing. True rejections: invented terms dropped; false rejections: expert terms dropped. *nameres*: the term has no candidate in the production Name Resolver (v1.7.0, Babel 2026jul22); *llm*: a model of another family (`llama3:8b`, batches of forty) does not recognise it; *union*: either. Measured on 6 October 2026 (`data/publication/gate_modes_v2_prod.json`). The two expert terms lost in union mode, *repeats* and *upper-arms*, were kept by the *llm* mode run alone: the second-family model's answer depends on the composition of the batch it receives, which differs once the resolver has removed its rejections.

| Gate mode | true rejections (of 34) | false rejections (of 32) |
|---|---|---|
| nameres | 19 | 0 |
| llm | 24 | 0 |
| union (published) | 32 | 2 |

Table S3: Typing accuracy on the 664 audited public mentions, every listed expert reading accepted (ambiguous terms included), overall and per coarse stratum of the Biolink class (reading scopes of the profiles). Macro: mean over the four strata.

| System | all (95% CI) | biochemical core | clinical core | shared | outside | macro |
|---|---|---|---|---|---|---|
| BiomedCAT, ReAct agent | 0.645 (±0.036) | 0.766 | 0.638 | 0.701 | 0.520 | 0.656 |
| BiomedCAT, single call | 0.610 (±0.037) | 0.759 | 0.594 | 0.624 | 0.505 | 0.620 |
| BiomedCAT, retriever top-1 | 0.381 (±0.037) | 0.517 | 0.356 | 0.401 | 0.287 | 0.390 |
| Claude Opus 5 | 0.687 (±0.035) | 0.876 | 0.525 | 0.771 | 0.614 | 0.696 |
| DeepSeek-V4-Flash | 0.633 (±0.037) | 0.759 | 0.588 | 0.713 | 0.515 | 0.644 |
| gemma-4-31b | 0.545 (±0.038) | 0.821 | 0.381 | 0.535 | 0.485 | 0.556 |
| mistral-small-24b | 0.449 (±0.038) | 0.386 | 0.494 | 0.554 | 0.376 | 0.453 |

#### 2.3 Audited benchmark: results by stratum

The audit also measures the generator: over the 664 mentions, 44.9% of the original labels were confirmed, 28.8% relabelled, 3.6% judged not to be an entity, 22.1% ambiguous and 0.6% (four mentions) naming a person; the error rate is lowest in the biochemical core (18.6% wrong) and highest in the clinical core and outside strata (40.0% and 42.6%), and higher for abbreviations (32.4% wrong, 30.0% ambiguous) than for canonical names (29.9%, 20.4%).

Table S4: Descriptions of the two slides per condition: wall-clock seconds, words, numbered items, [UNCLEAR] marks, and elements of the slide detected by the automatic token check (slide 1: 24 elements; slide 2: 21). Item counts for Qwen include indented sub-bullets.

| | | | Slide 1 | | | | | Slide 2 | | |
|---|---|---|---|---|---|---|---|---|---|---|
| Model, $T$ (seed) | Profile | s | words | items | uncl. | elem. | s | words | items | uncl. | elem. |
| Gemma, 0 | none | 19 | 222 | 8 | 0 | 18 | 19 | 276 | 10 | 0 | 15 |
| Gemma, 0 | biochemical | 19 | 275 | 10 | 0 | 17 | 17 | 199 | 8 | 0 | 11 |
| Gemma, 0 | genetic | 19 | 260 | 9 | 0 | 18 | 22 | 315 | 13 | 0 | 14 |
| Gemma, 1 (0) | none | 21 | 277 | 7 | 2 | 20 | 17 | 214 | 5 | 2 | 14 |
| Gemma, 1 (0) | biochemical | 18 | 225 | 7 | 0 | 17 | 20 | 300 | 9 | 1 | 14 |
| Gemma, 1 (0) | genetic | 21 | 278 | 6 | 5 | 16 | 20 | 233 | 7 | 0 | 13 |
| Gemma, 1 (1) | none | 19 | 244 | 8 | 0 | 21 | 19 | 254 | 7 | 1 | 13 |
| Gemma, 1 (1) | biochemical | 18 | 222 | 6 | 1 | 18 | 19 | 245 | 8 | 1 | 13 |
| Gemma, 1 (1) | genetic | 19 | 229 | 7 | 0 | 18 | 21 | 267 | 9 | 1 | 12 |
| Qwen, 0 | none | 109 | 229 | 9 | 0 | 20 | 110 | 288 | 14 | 0 | 10 |
| Qwen, 0 | biochemical | 110 | 257 | 12 | 0 | 21 | 109 | 277 | 10 | 0 | 13 |
| Qwen, 0 | genetic | 113 | 279 | 15 | 0 | 21 | 110 | 215 | 10 | 0 | 11 |

**Table S6.** Experimental log for the selection of the typing configuration. **Obs.**: observation; **Hyp.**: hypothesis. **Layer**: workflow layer on which the change acts (retriever, agent reasoning, or model parameter). **Result**: hit@1 for retriever experiments (1,075 public terms) and accuracy for agent experiments (300 reference terms, original gold), unless stated otherwise. Differences are given relative to the configuration current at the time of the experiment.

| ID | Layer | Observation and hypothesis | Change tested | Result | Conclusion |
|---|---|---|---|---|---|
| E1 | Retriever | **Obs.** Biolink definitions are semantically far from the queried text. **Hyp.** Exemplars more similar to the retrieved text improve retrieval. | **Exemplars (v3 → v4).** A leg of synthetic exemplar mentions added to the reciprocal rank fusion. | Retriever hit@1 +7 points. Agent without exemplars: 0.373 (0.410 with). | Improvement above 5 points: hypothesis validated, exemplars kept. |
| E2 | Retriever | **Obs.** Synthetic benchmarks can be far from real use. **Hyp.** Names and synonyms taken from the knowledge graph improve retrieval. | **Index v5.** Synthetic exemplars replaced by real KG2c node names (52 classes). | Retriever hit@1 −2 points. | No improvement, variant abandoned. KG2c names stay close to identifiers that scientists rarely use in their prose. |
| E3 | Retriever (candidate list) | **Obs.** A quick test identified the number of retrieved candidates $N$ as a parameter with a large effect. **Hyp.** Increasing $N$ to 15 improves accuracy by more than 5 points. | **15 candidates.** The agent search tool returns 15 classes instead of 10. | 0.443 (+3.3). On the full 1,356 terms: 0.455 against 0.4565. | No significant improvement. Retriever hit@15 rises but the agent decision does not change at scale. |
| E4 | Agent reasoning | **Obs.** Biolink definitions can differ from those of other ontologies, creating a gap between what the model knows and what it is asked to do. **Hyp.** Imposing the hierarchy in the prompt keeps the decision within the Biolink structure. | **Tree skeleton.** Biolink hierarchy (class names only) added to the system prompt. | 0.410 (no change). | No effect. The Gemma models accept a long context, but filling it degrades the reasoning of a small model. |
| E5 | Retriever (branch scores) | **Obs.** The agent often chose a class outside the correct Biolink branch. **Hyp.** Aggregating the evidence of a whole branch first guides the agent to the right branch. | **Aggregation by branch, smoothing.** Retriever scores cascaded down the branches, with parent-based smoothing. | Negative. | No improvement and a longer process, likely because of the added prompt information. |
| E6 | Model parameter | **Obs.** Larger models usually hold more knowledge, and a 12-billion-parameter model still fits (at the limit) a 16 GB RAM set-up. **Hyp.** A larger local model improves accuracy. | **gemma4:12b-it-qat** instead of `e4b-it-qat`, same prompt. | 0.387 (−2.3). | Within the same family, a larger local model does not improve reasoning. |

*Continued on next page*

| ID | Layer | Observation and hypothesis | Change tested | Result | Conclusion |
|---|---|---|---|---|---|
| E7 | Model parameter | **Obs.** The Gemma 4 documentation recommends higher temperatures given its architecture, warning of degraded output at low temperature, whereas temperature 0 had been chosen to limit hallucination. **Hyp.** Sampling at the recommended temperature stabilizes Gemma 4. | **T = 1.** Recommended sampling (temperature 1, top-k 64, top-p 0.95) instead of greedy decoding. | Agent 0.433 (+2.3). Real FSHD document: exact recall −25 and −31 points in two runs; the two runs were not identical. | Gain below threshold on isolated terms and a large loss of recall on the document: sampling rejected, T = 0 kept. |
| E8 | Agent reasoning | **Obs.** Some candidates are hard to resolve in three steps. **Hyp.** A second round of reasoning improves accuracy. | **Second-pass iterations.** Hard terms replayed with 20 candidates and 5 steps; a variant adds hidden reasoning. | 0.423 / 0.427. | No significant improvement: the model most often reached the same conclusion. |
| E9 | Agent reasoning | **Obs.** Local models run with limited memory, which restricts the usable context. **Hyp.** A telegraphic style shortens the reasoning and improves accuracy. | **Telegraphic thought.** Reasoning lines limited to a few words. | 0.317 (−9.3). | No improvement; the reasoning became longer, not shorter. |
| E10 | Agent reasoning | **Obs.** The two passes of E8 reach the same conclusions. **Hyp.** A reasoning trace prevents the model from falling into the same trap twice. | **Reasoning in the loop.** Hidden reasoning (1,500 tokens) activated at the first pass. | 0.430 (+2). | Below threshold in the loop. Hidden reasoning was kept for the single-call mode, where it is measured together with the menu (E14). |
| E11 | Retriever (sparse leg) | **Obs.** Accuracy is mainly limited by retriever hit@10. **Hyp.** A sparse leg tokenized with a biomedical vocabulary improves ranking. | **BM25.** Okapi BM25 leg, tokenized with scispaCy, added to the fusion. | Hit@1 0.334 against 0.333. Same result (no gain) on the 997-mention KG2c benchmark. | No improvement. The scispaCy model is slow to load at application start; the leg is optional and off by default. |

*Continued on next page*

| ID | Layer | Observation and hypothesis | Change tested | Result | Conclusion |
|---|---|---|---|---|---|
| E11b | Retriever (dense encoder) | **Obs.** As E11. **Hyp.** A dense encoder specialized in biomedical language improves ranking over the general-purpose encoder. | **Encoder comparison.** Five alternative sentence encoders substituted for the default one, including SapBERT and `pubmedbert-base-embeddings`. | Hit@1 on the 1,075 public terms, default encoder (`bioclinical-modernbert-base-embeddings`) 0.333: `pubmedbert-base-embeddings` 0.348 (+1.5); `all-mpnet-base-v2` 0.314 (−2.0); `S-PubMedBert-MS-MARCO` 0.303 (−3.0); `bge-large-en-v1.5` 0.300 (−3.4); SapBERT 0.294 (−3.9). | No encoder above threshold; the default encoder was kept. Specialized encoders also add load time at application start. |
| E12 | Retriever (lexical legs) | **Obs.** Gene names and other acronyms were hard to retrieve. **Hyp.** An exact keyword search over real acronyms improves retrieval. | **Exact lexical legs.** Exact match on class names and aliases, then on exemplars and KG2c synonyms. | Hit@1 +0.5 (synthetic); +12 (mentions of the real document, 0.16 → 0.28). Agent 0.407 against 0.410. | Large gain on real mentions, neutral on synthetic terms; agent unchanged. Kept at weight 1 at the time; switched off in the published runs (`RAG_LEXICAL_WEIGHT=0`, `RAG_KG2C_LEXICAL=0`), whose retriever uses the dense and exemplar legs only. |
| E13 | Agent reasoning | **Obs.** The ReAct loop is slow on local hardware. **Hyp.** Prompt design alone can save time while keeping accuracy. | **Decide mode, D10.** No loop: a single call per term over the ten retriever candidates, with hidden reasoning. | 0.427 (+1.7). | Below threshold and no time gain: hidden reasoning costs 18–24 s per term against about 15 s for the loop. |

*Continued on next page*

| ID | Layer | Observation and hypothesis | Change tested | Result | Conclusion |
|---|---|---|---|---|---|
| E14 | Agent reasoning | **Obs.** As E13. **Hyp.** Giving the model the local Biolink hierarchy lets it reason on the hierarchy. | **Decide mode, D10N.** Menu extended with children, siblings and parent of the first three candidates (maximum 30). | 0.467 (+4.0 over D10, +5.7 over the tool loop); 0.470 once integrated with the raw prompt (E19). With 20 candidates and 40 menu entries (D20N): 0.430. | Best configuration of the series. The gain over D10 is below threshold and a longer menu (D20N) loses it: prompt length degrades the model. |
| E15 | Agent reasoning | **Obs.** Prompt length degrades the model (E14). **Hyp.** A restricted amount of hierarchical knowledge is enough to reason on the hierarchy. | **Neighbour grid.** Neighbours of the first candidate (N1); neighbours of the first three plus mixins (N3M); neighbours of the first five (N5). | N1 0.423; N3M 0.460; N5 0.437. | No significant improvement. Mixins (horizontal links across the Biolink hierarchy) do not help. |
| E16 | Agent reasoning | **Obs.** Some class names are repetitive and bias the prompt vocabulary towards one Biolink parent. **Hyp.** Removing grouping classes from the menu reduces this bias. | **Excluded grouping parents.** Classes of the form “X or Y” removed from the added neighbours. | 0.470 (no change); four additional `NONE` verdicts. | No significant improvement. |
| E17 | Agent reasoning | **Obs.** Menu definitions were cut at 150 characters, in the middle of a sentence for about 30% of them. **Hyp.** Longer definitions improve reasoning. | **Definitions at 250 characters.** Cut at sentence boundaries instead of a raw 150-character limit. | 0.417 (−5.3). | Worse. More definition text does not help: the beginning of a definition is its most informative part. |
| E18 | Agent reasoning | **Obs.** Detecting the correct branch is still the bottleneck. **Hyp.** A provenance line for each retrieved class and longer reasoning help when the choice is ambiguous. | **Prompt v2.** Menu grouped by parent, provenance line, four steps of private reasoning. | 0.430 (+1.3 on the same slice as the 250-character baseline of E17). | No significant improvement. |
| E19 | Agent reasoning | **Obs.** A role label (“User:”) was still present in the prompt. **Hyp.** Removing it shortens the context and changes how the model reads the task. | **“User:” prefix.** Same prompt sent with and without the role label. | Raw prompt 0.470; labelled prompt 0.413. | Significant (5.7 points): a role label alone changes the model’s perspective on the task. Raw prompt kept. |

*Continued on next page*

| ID | Layer | Observation and hypothesis | Change tested | Result | Conclusion |
|---|---|---|---|---|---|
| E20 | Agent reasoning (runtime) | **Obs.** Unloading the model after each call, used to save memory, costs speed and prevents caching. **Hyp.** Keeping the model resident improves speed and verdict consistency. | **Resident model.** Model kept loaded for 10 minutes (`keep_alive`) instead of being unloaded after each call; identical verdicts between two runs as the measure. | 50 public terms, machine otherwise idle: three runs with the model unloaded after each call (two with seed 0, one without seed) gave 100% identical verdicts; two back-to-back runs with the model resident agreed on 82%, and on 70 to 74% with the unloaded runs. Complete tool-loop runs on the 1,356 terms made with a resident model or while other experiments shared the machine (5, 10 and 11 Sept): 69 to 72% identical. Two complete single-call runs, model unloaded after each call: 99.7% identical. | Residency lowers reproducibility instead of raising it. The tool loop is reproducible when the model is unloaded after each call on an idle machine; its variation comes from the serving conditions (resident model, concurrent jobs). The model is unloaded after each call (`OLLAMA_KEEP_ALIVE=0`) in the published configuration. |

#### 2.4 Multi-scale case

### 3 Supplementary Figures

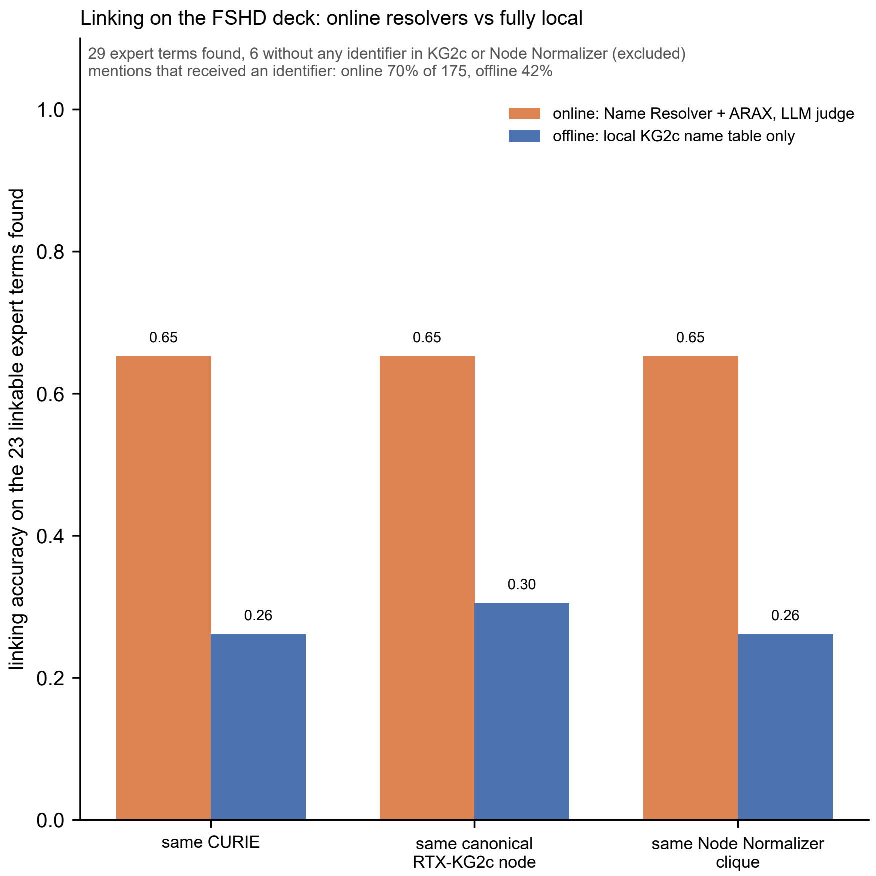


Figure S1: **Entity linking on the FSHD deck, online resolvers against the fully local mode.** Reading and typing are identical in both modes (same 175 mentions; extraction recall 0.91 on the 32 explicit expert terms and typing accuracy 0.48 on the 29 found ones); only the linking stage differs. Online: candidate identifiers from the Name Resolver (unconstrained and constrained to the predicted Biolink class) and ARAX, canonicalised to RTX-KG2c, selected by the local judge model; offline: exact name and synonym match against the local RTX-KG2c name table, no term leaves the machine. Accuracy on the 23 found expert terms that have an identifier in RTX-KG2c or Node Normalizer (6 of the 29 found terms have none and are excluded: D4Z4 array, repeats, impaired myogenesis, DNA binding domains, DUX4 binding site, DUX4 target gene), under three nested criteria: same CURIE string; same canonical RTX-KG2c node; same Node Normalizer clique (conflation off). The clique criterion is applied by the evaluation to both modes; the offline pipeline itself never calls Node Normalizer. Mentions that received any identifier, right or wrong: online 70% of 175, offline 42%.

Table S5: Elements printed or drawn on the two slides and their detection in each condition (automatic, case-insensitive token match; `x` = present). Columns: G0 = Gemma $T = 0$; G1a, G1b = Gemma $T = 1$, seeds 0 and 1; Q0 = Qwen $T = 0$; n/b/g = no profile / biochemical / genetic.

| | G0 | | | G1a | | | G1b | | | Q0 | | |
|---|---|---|---|---|---|---|---|---|---|---|---|---|
| Element | n | b | g | n | b | g | n | b | g | n | b | g |
| *Slide 1* | | | | | | | | | | | | |
| FSHD / facioscapulohumeral | x | x | x | x | x | x | x | x | x | x | x | x |
| muscular dystrophy | x | x | x | x | x | x | x | x | x | x | x | x |
| no treatment / cure | | | | | | x | x | x | | x | x | x |
| face | x | x | x | x | x | x | x | | x | x | x | x |
| shoulders | x | x | x | x | x | | x | | | x | x | x |
| upper arms | x | x | x | x | x | | x | | x | x | x | x |
| hypomethylation | x | x | x | x | x | x | x | x | x | x | x | x |
| D4Z4 (as written) | | | | | | | | | | x | x | x |
| chromosome 4 | x | x | x | x | x | x | x | x | x | x | x | x |
| toxic DUX4 protein | x | | | x | x | x | x | x | x | x | x | x |
| DUX4 | x | x | x | x | | x | x | x | x | x | x | x |
| FSHD1 (as written) | | | x | x | | | x | x | x | x | x | x |
| contraction of the array | x | x | x | | x | x | x | x | x | x | x | x |
| FSHD2 (as written) | | | x | x | | | x | x | x | x | x | x |
| SMCHD1 | x | x | x | x | x | x | x | x | x | x | x | x |
| DNMT3B | x | x | x | x | x | | x | x | x | x | x | x |
| epigenetic modifier genes | x | x | x | x | x | x | x | x | x | x | x | x |
| reduced methylation | x | x | x | x | x | x | x | x | x | x | x | x |
| exon 1 | | x | | x | x | x | x | x | | | | |
| exon 2 | x | x | x | x | x | x | x | x | x | x | x | x |
| unstable DUX4 mRNA | x | x | x | x | x | x | x | x | x | x | x | x |
| stable DUX4 mRNA | x | x | x | x | x | x | x | x | x | | x | x |
| poly(A) tail, “(A)n” | x | | | x | | | | | | | | |
| reference line | | | | | | | | | | | | |
| *Slide 2* | | | | | | | | | | | | |
| DUX4 | x | x | x | x | x | x | x | x | | x | x | x |
| FSHD | x | x | x | x | x | x | x | x | x | x | x | x |
| DNA binding domains | x | | x | x | | | | | x | | x | |
| HD1 / HD2 | x | x | x | x | x | | x | x | | | x | x |
| N-ter / C-ter | x | x | x | x | x | x | x | x | x | | | x |
| TAD | x | | x | x | x | x | x | x | x | | | |
| binding-site motif | | | | | | | | | | | | |
| DUX4 binding site | | | | | x | x | x | | | x | x | |
| DUX4 target gene | x | | | x | | | | | | | x | |
| protein aggregation | x | x | x | x | x | x | x | x | x | x | x | x |
| impaired NMD | | x | x | x | x | x | x | x | x | x | x | x |
| ROS / oxidative stress | x | x | x | x | x | x | x | x | x | x | x | x |
| mitochondrion | | | x | | | | | | | | | |
| impaired myogenesis | x | x | | x | x | x | x | x | x | | x | |
| cell death | x | x | x | x | x | x | x | x | x | x | x | x |
| toxicity | x | x | x | x | x | x | | x | x | x | x | x |
| damage | | | x | | x | x | | x | x | x | x | x |
| control vs FSHD histology | x | x | x | x | x | x | x | x | x | x | | x |
| inflammatory infiltrate | x | | | | | | x | | | | | |
| fluorescence panels | | | | | | | | | | | | |
| reference line | x | | | | | | | | | | | |

Table S7: Pathway signature nodes (RTX-KG2c names). Reactome entries carry the suffix *(Homo sapiens)* in the graph.

| Signature | Nodes |
|---|---|
| A, innate antiviral sensing | DDX58/IFIH1-mediated induction of interferon-$\alpha/\beta$; RIG-I signaling pathway; Toll Like Receptor 3 (TLR3) Cascade; ISG15 antiviral mechanism; Activation of IRF3, IRF7 mediated by TBK1, IKK$\epsilon$; TRAF6 mediated IRF7 activation; antiviral innate immune response (GO); cGAS/STING signaling pathway; antigen processing and presentation of peptide antigen via MHC class I (GO); cellular response to dsRNA (GO) |
| B, mitochondria and redox | Mitochondrial complex I (NADH dehydrogenase); reactive oxygen species metabolic process (GO); Aerobic respiration and respiratory electron transport; The NLRP3 inflammasome; NLRP3 (gene); HMGB1 (gene); 4-hydroxynonenal; cell death in response to oxidative stress (GO); Interleukin-6 family signaling; Detoxification of Reactive Oxygen Species |
| C, remodelling and fibrosis | Extracellular matrix organization; Degradation of the extracellular matrix; collagen-containing extracellular matrix (GO CC); Signaling by TGF-beta Receptor Complex; TGFB1 (gene); Chemokine CCL2; TNF-$\alpha$ (protein); Tumor Necrosis Factors; execution phase of apoptosis (GO); Collagen |

Table S8: Structure of the kept graphs. Pairs: seed pairs connected inside the kept graph. Mean distance: mean shortest path between connected seed pairs (hops). Saturated: Biolink categories at the per-category cap of 300.

| Slide | Seeds | Nodes | Edges | Diameter | Components | Pairs | Mean dist. | Saturated |
|---|---|---|---|---|---|---|---|---|
| A, molecular | 20 | 5,491 | 51,748 | 13 | 1 | 190/190 | 3.43 | 14/31 |
| B, organelle | 19 | 5,500 | 52,303 | 13 | 2 | 153/171 | 3.54 | 15/32 |
| C, tissue | 24 | 5,604 | 53,136 | 15 | 3 | 231/276 | 3.06 | 15/32 |
| A+B+C, integrated | 20 | 6,080 | 74,541 | 14 | 2 | 171/190 | 3.97 | 18/31 |
| Summary slide | 25 | 5,959 | 112,475 | 16 | 5 | 210/300 | 3.85 | 15/35 |

Table S9: Recovery of the pathway signatures in each kept graph: nodes present out of ten, and in brackets the median rank percentile of the best hits inside their Biolink categories (0 = top).

| Signature | A | B | C | A+B+C | Summary |
|---|---|---|---|---|---|
| A, innate antiviral | 9 (0.16) | 0 | 4 (0.45) | 6 (0.63) | 3 (0.71) |
| B, mitochondria and redox | 1 (0.46) | 5 (0.06) | 6 (0.55) | 5 (0.68) | 6 (0.45) |
| C, remodelling, fibrosis | 3 (0.58) | 3 (0.35) | 8 (0.07) | 9 (0.13) | 7 (0.33) |

Table S10: Minimum number of hops inside each kept graph between the three signatures and from DUX4 to each signature (undirected; "–": no path, the signature being absent from the graph). In brackets, the length of the best-weighted path when it differs.

| Slide | A–B | A–C | B–C | DUX4–A | DUX4–B | DUX4–C |
|---|---|---|---|---|---|---|
| A | 2 | 3 | 4 | 3 | 4 | 2 |
| B | – | – | 2 | – | 1 | 3 |
| C | 4 (5) | 3 | 2 (3) | 4 | 1 | 3 (4) |
| A+B+C | 2 | 2 | 2 (3) | 3 | 2 | 3 (4) |
| Summary | 2 | 2 | 2 (3) | 4 | 2 | 3 |
